# Intra-individual changes to peak alpha frequency map to pain perception following high frequency sensitisation of central nociceptive pathways: A test-retest reliability study

**DOI:** 10.64898/2026.10.31.763331

**Authors:** Samuel Mugglestone, Sam Hughes, Giorgio Ganis

## Abstract

Resting-state EEG can capture cortical dynamics relevant to sensitised pain states in the absence of ongoing nociceptive engagement. However, whilst the slowing of global oscillations is evident within various chronic pain conditions, the mechanistic specificity of PAF and α-band power remains uncertain. Therefore, the present study assessed these features across two identical test-retest sessions in which HFS was applied to induce the development of SHA as characterised through 128 mN MPS. Crucially, changes in α-band features post-HFS were investigated. Blind source separation was also applied to identify independent sensorimotor components relevant to nociception and assess their test-retest reliability. Results demonstrated good test-retest reliability for sensor-level data at baseline, with inter-individual changes in global PAF reflecting MPS post-HFS. However, no specific regional driver of this signal was evident. Alternatively, components identified through sensorimotor template matching revealed a significant influence of α-band power on MPS both at baseline and post-HFS. However, only a small number of these components genuinely localised to pre- or post-central gyri, and test-retest reliability of α-band power was poor. Together, these findings indicate that resting-state EEG may be useful for tracking changes in cortical α-band oscillations alongside the development of pain sensitivity. However, methods of sensorimotor component identification evidenced limited agreement and global PAF slowing could not be linked to a specific electrode grouping.

## Introduction

Resting-state electroencephalography (EEG) characterises neural dynamics separate to task-related cortical processing (Perinelli & Ricci, 2025), allowing for the investigation of altered neural states related to pain sensitivity in the absence of ongoing nociception (Ploner et al., 2017). Within chronic pain cohorts, abnormal oscillations have been identified across a wide variety of conditions (Ahmad & Barkana, 2025; Cherin et al., 2026; González-Villar et al., 2017; Mussigmann et al., 2022; Ploner et al., 2017). However, the specificity of these neurological markers varies across cohorts (Ahmad & Barkana, 2025; Mussigmann et al., 2022; Ploner et al., 2017), indicating the necessity of mechanistically-focused investigations to assess the development of maladaptive pain processing (Bannister & Hughes, 2023). However, whilst altered resting-state EEG dynamics have been characterised within sensitised pain states, the development time course of induced changes requires further investigation, particularly concerning test-retest reliability across repeated experimental sessions (Chowdhury et al., 2025; Furman et al., 2020; Lebrun, Lenoir, et al., 2025; Lebrun, Ricci, et al., 2025; Leone et al., 2025).

The present study applied high-frequency stimulation (HFS) to induce spinal cord dorsal horn plasticity, manifesting as cutaneous secondary hyperalgesia (SHA) as early as 15 minutes post-HFS, which can be maintained over hours (Fawsitt-Jones et al., 2024; Klein et al., 2004; Klein et al., 2008; Lebrun, Lenoir, et al., 2025; Leone et al., 2025; Pfau et al., 2011; Quesada et al., 2021; van den Broeke et al., 2015). We have previously demonstrated that variations in the time course of SHA development can occur and are not directly attributable to typically considered psychophysical factors but may be captured by certain aspects of nociceptive-evoked cortical activity (Mugglestone, Ganis, et al., 2026; Mugglestone, Hughes, et al., 2026). Now, using the same behavioural dataset, we investigate the reliability and utility of resting-state EEG dynamics for assessing altered states of cortical activity that may underly variability in the development of SHA. Notably, HFS sensitisation is not accompanied by an ongoing tonic pain sensation, and therefore, altered cortical dynamics may be experimentally isolated to an altered sensitivity state.

Of particular interest is the slowing of the peak alpha frequency (PAF) which is implicated within many long-term chronic conditions (Cavaleri et al., 2025; Cherin et al., 2026; Fauchon et al., 2022; Liberati, 2025; Mussigmann et al., 2022; Ploner et al., 2017) and reflects alterations to corticothalamic feed-back signalling originating within the infragranular cortex (El Zghir et al., 2024; Michalareas et al., 2016; Ploner et al., 2017). Recent investigations within chronic pain populations have demonstrated that slowed PAF is often present, potentially developing in response to widespread persistent pain (Cavaleri et al., 2025; Cherin et al., 2026; Fauchon et al., 2022; Mussigmann et al., 2022). However, the mechanistic specificity of PAF remains in question, as many chronic pain subtypes do not exhibit significant changes (Ahmad & Barkana, 2025; McLain et al., 2022; Mussigmann et al., 2022). Accordingly, experimental assessments of induced pain have demonstrated that PAF is unchanged by brief noxious stimuli (May et al., 2025), whilst substantial alterations are evident during tonic pain respective to pain intensity (Chowdhury et al., 2025; Furman et al., 2018; Furman et al., 2020; Furman et al., 2019; Nir et al., 2010). Therefore, altered PAF may simply result from ongoing nociception, although this does not account for chronic pain states that do not demonstrate a persistent painful experience. Furthermore, one investigation has applied HFS to assess the role of PAF in capturing the induction of altered sensitivity states, but have demonstrated no significant relationship between baseline PAF prior to HFS conditioning and the magnitude of post-HFS SHA (Lebrun, Ricci, et al., 2025). Additionally, this result has been mirrored by clinical evidence demonstrating that pre-operative PAF is predictive of acute but not chronic postoperative pain (Han et al., 2025). Therefore, rather than representing a pre-existing trait of vulnerability to sensitivity development, altered PAF may emerge alongside the development of altered states of nociceptive processing. Accordingly, understanding how PAF alterations reflect ongoing neurophysiological changes during sensitivity development is crucial. However, whilst Lebrun, Ricci, et al. (2025) report data on the predictive utility of baseline PAF, they do not consider whether PAF was altered by HFS-induction throughout the time course of SHA development. Therefore, it is critical to provide data on the reliability, sensitivity, and specificity of PAF changes post-HFS.

The altered PAF present during states of chronic pain can be effectively characterised at the sensor-level by the global average across electrodes (Cavaleri et al., 2025; Cherin et al., 2026; McLain et al., 2022), potentially reflecting widespread alterations to postsynaptic communication within thalamocortical circuits (Da Silva & Van Leeuwen, 1977; Da Silva et al., 1980; El Zghir et al., 2024; Halgren et al., 2019; Michalareas et al., 2016; Ploner et al., 2017). Therefore, within these processes may be related to the development of altered states of pain sensitivity. However, sensor-level data cannot facilitate the identification of specific underlying cortical mechanisms from which altered PAF arises. Therefore, recent investigations within experimentally induced pain paradigms have focused upon the identification of relevant ICs using blind source separation to capture unique variance attributed to a single representative source (Chowdhury et al., 2025; Delorme & Makeig, 2004; Delorme et al., 2012; Hsu et al., 2018; Lebrun, Ricci, et al., 2025; Palmer et al., 2012; Viola et al., 2009), typically termed the sensorimotor component due to a characteristic topography centred above pre- and post-central gyri (Cherin et al., 2026; Chowdhury et al., 2025; Furman et al., 2018; Furman et al., 2020, 2021; Furman et al., 2019; Lebrun, Ricci, et al., 2025; Liberati, 2025; May et al., 2025).

However, the consistent identification of this sensorimotor component across studies and populations remains a pressing concern, especially as attempts to replicate methods within patient cohorts has been met with limited success (Cherin et al., 2026). Furthermore, it is unknown whether the sensorimotor components identified by previous research reflect genuine and specific sensorimotor engagement or are the result of co-occurring cortical processes across multiple disparate regions. Primarily, a correlation-based template matching approach has been used to identify sensorimotor components across individuals (Cherin et al., 2026; Chowdhury et al., 2025; Furman et al., 2018; Furman et al., 2020, 2021; Furman et al., 2019). However, this is dependent upon accurate specification of a template component and may not generalise across EEG recording setups. Comparatively, the use of study-level component clustering through the application of optimised K-means has shown success but has yet to be applied to pain-related PAF identification (Delorme & Makeig, 2004; Handiru et al., 2024; Kim et al., 2022; Miyakoshi et al., 2021; Rousseeuw, 1987; Thammasan & Miyakoshi, 2020). Alternatively, given the nomenclature specifying sensorimotor regions of the cortex, a source-based identification of components localised to pre- or post-central gyri may be most appropriate (Desikan et al., 2006).

Therefore, in the present study, the test-retest reliability of resting-state EEG dynamics across multiple time points following HFS conditioning was investigated. It was hypothesised that within both sessions, changes in PAF assessed both at the global level and at central electrodes would reflect MPS throughout the SHA development time course. It was also anticipated that sensorimotor components identified through templated matching would capture this signal and localise to pre- and post-central gyri indicating altered oscillatory activity within sensorimotor cortices following HFS.

## Methods

### Pre-registration

This study is part of a wider research program investigating the reliability of psychophysical and neurophysiological methods for assessing temporal changes to central nociceptive function induced by HFS conditioning. The overall study protocol was preregistered on the Open Science Framework (OSF) and is available at: https://doi.org/10.17605/OSF.IO/S5A9Y

### Participants

Recruitment was conducted via the University of Plymouth’s Paid Participation Pool alongside word of mouth. Participants were instructed to abstain from alcohol and drugs prior to each visit, and to specifically avoid taking painkillers and to limit caffeine intake. At the start of each visit compliance was confirmed. All 35 recruited participants (final sample size = 27) provided written informed consent and were informed of their continual right to withdraw. The study was approved by the University of Plymouth Ethics Committee (ID:3777) and in compliance with the Declaration of Helsinki (Association, 2013).

Exclusion criteria resulted in the removal of one participant: no history of neurological conditions, long-term health conditions, chronic pain, psychiatric conditions, steroid use, inflammatory disease, or immunosuppression; no allergies or severe skin conditions that may affect the methods or areas of testing; not currently pregnant.

### Experimental design

Two functionally identical sessions were conducted at least a week apart with the aim of assessing the between-session reliability of the temporal changes in EEG features following HFS-conditioning. Various behavioural data were collected at the beginning of and throughout each session which are not relevant to the present publication and are discussed separately (Mugglestone, Ganis, et al., 2026). Prior to beginning the main experimental procedures, an EEG electrode array was set up and tested for impedance, before participants were familiarised with the sensory testing stimuli and electrical detection threshold (EDT) was determined over the volar surface of the forearm. Baseline mechanical pinprick sensitivity (MPS) responses were assessed on the ipsilateral and contralateral forearms, followed by HFS conditioning. MPS responses were then reassessed at 4 time points post-HFS with 15-minute intervals between the beginning of each block. During stimulus delivery, view of the testing sites was occluded via a screen-box which limited the perceptual field to a computer monitor on which a fixation cross (white) was displayed throughout. Participants were instructed to focus their visual attention onto this fixation cross and to refrain from blinking or moving as much as possible. A chinrest was also provided to reduce muscle-strain artifacts.

Within each block, 20 stimuli were delivered per condition (arm) with an interstimulus interval (ISI) of ∼2 seconds resulting in 40 stimuli per arm, and a total of 40 MPS stimuli within each time point. This was considered the minimum number of trials to effectively characterise event-related EEG features based upon comparable studies (Biurrun Manresa et al., 2018; Gousset et al., 2023; van den Broeke et al., 2020; van den Broeke et al., 2017; Van den Broeke et al., 2016; van den Broeke et al., 2015).

### High-Frequency Stimulation

Cutaneous electrical stimuli were applied to the right volar forearm (5cm distal to the cubital fossa) using an EPS-P10 electrode (MRC Systems GmbH, Heidelberg, Germany) via a constant current stimulator (2ms pulse width; DS7; Digitimer Ltd; Welwyn Garden City, UK). Prior to attaching the HFS electrode, the skin was cleaned with a non-alcoholic aqueous solution and allowed to dry. The EPS-P10 electrode contains a circular array of 10 cathodal pins (diameter: 0.25mm) and a larger surface area anode (area: 410 mm2). Such concentric surface electrodes have been shown to selectively activate A-delta and C fibre free nerve endings located in the superficial epidermal layers without the excitation of A-beta fibre mechanoreceptors located in the dermis (Medina & Hughes, 2025; Poulsen et al., 2022). Sufficiently high intensity electrical stimuli can bypass peripheral transduction mechanisms directly activating voltage-gated ion channels (Leone et al., 2025). Notably, this specificity may falter depending upon the intensity of stimulation, resulting in the activation of deeper A-beta fibres (Legrain & Mouraux, 2013). However, characterising the incidence of this phenomenon is beyond the scope of the current study.

Electrical detection threshold was determined using 2ms pulses starting at 0.05 mA in 0.05 mA increments until participants reported a clear sensation, then decreased by 0.01 mA until the sensation was lost. The intensity was increased in 0.01 mA increments until the sensation returned, repeating this process three times. The average of the six recorded intensities represented each participant’s EDT. Trains of HFS (100 Hz; NL304 Period Generator; Digitimer Ltd) were applied 5 times to the test (right arm) for 1 second each (NL405 Width Delay; Digitimer Ltd) at 15x EDT every 10 seconds (i.e. over a 50 second period). The participants were asked to rate the intensity of the HFS after each train using a numerical rating scale (NRS; 0 = no pain, 100 = worst pain imaginable).

### Heterotopic mechanical stimulation

Mechanical pain sensitivity (MPS) was assessed across both forearms using a 128 mN pin prick stimulator within a typical heterotopic area laterally adjacent to the conditioning electrode yet within 2 cm as depicted in Lebrun, Lenoir, et al. (2025) and Medina and Hughes (2025). However, the area of SHA was not systematically mapped. Although mechanical pinprick stimuli activate both high- and low-threshold mechanoreceptors, low-intensity pinpricks can be perceived as non-painful (Leone et al., 2021). Therefore, the medium range intensity of 128mN was used throughout the study and baseline pain ratings are reported.

Following every five successive stimuli, participants rated their average perceived pain on a numerical rating scale (NRS; 0 - 100). These were further averaged to provide a single value for each time point. Therefore, within each time point, participants received a total of 20 pinprick stimuli to each of the ipsilateral and contralateral arms.

### EEG data recording

EEG data were recorded continuously throughout the main experimental procedures using an ActiCAP Slim 64-channel electrode array positioned according to the extended 10-20 system, referenced to FCz and grounded at AFz (band-pass filter = 0.01 to 100 Hz; sampling rate = 500 Hz) using an ActiCHamp Plus amplifier (Brain Products GmbH, Glitching-Sȕd, Germany).

### EEG data preprocessing

Preprocessing was conducted using EEGLAB (version 2025.1.0) and custom functions within MATLAB (version 2024b) as follows (Delorme & Makeig, 2004):

- Continuous EEG data were band-pass filtered (0.1-100 Hz) using a zero-phase, non-causal, Hamming-windowed FIR filter (transition width = 0.1 Hz; −6 dB cutoff definition).
- Aberrant channels were identified and removed automatically using the Clean_rawdata plugin (version 2.10) without ASR or window rejection (flatline >= 5 seconds; channel correlation = 0.8; line noise = 4 std; high-pass filter for channel correlation = 0.25 Hz; 0.75 Hz). Channels were also manually inspected and removed when deemed appropriate (Delorme, 2023).
- UK mains interference (50 Hz and its first harmonic at 100 Hz) was removed using the CleanLine plugin (version 2.1).
- Aberrant data drifts were identified automatically using the Clean_rawdata plugin without channel or window rejection and artifact subspace reconstruction was applied (maximum acceptable 0.5 second, window deviation = 20 std) (Chang et al., 2018; Delorme, 2023).
- Data were re-referenced to average with care to avoid rank-deficiency issues (Delorme & Makeig, 2023; Kim et al., 2023).
- Independent component analysis (ICA) was conducted using AMICA (version 17) (Hsu et al., 2018), using a single model with 3000 iterations and all other parameters at default.
- Artifact components classified as muscle, heart, or channel noise were rejected with a threshold of ≥ 0.9 using the ICLabel plugin. Eye-related components were handled separately: components were first labelled “ocular” when the eye class had the highest ICLabel probability (i.e., was the dominant class). Among these ocular-labelled components, only those with a brain-class probability < 5% were removed. This approach targeted ocular artifacts whilst reducing the risk of discarding components with a non-trivial neural contribution, as done in previous work (Hsu et al., 2024).
- Resting-state data were segmented into half-overlapping epochs of 5 seconds within the 240 second eyes-closed window prior to each QST block, resulting in 95 epochs per time point (Furman et al., 2020; Lebrun, Lenoir, et al., 2025).
- Spherical interpolation of removed electrodes was applied to ensure comparability across datasets.

Notably, epoching and interpolation were conducted post-ICA to ensure that a continuous and valid data context was preserved, maximising AMICA performance. In total, 92% of channels were retained with an average of 4.6 (sd = 3.5) removed per dataset. Alternatively, ICLabel component rejection resulted in the retention of 75% of independent components (ICs), with an average of 5.6 (sd = 2.5) removed per dataset.

### Source Localisation

The source localisation of ICs was conducted using the DIPFIT plugin (version 5.6) through the estimation of equivalent-dipole locations as follows (Oostenveld et al., 2003; Oostenveld et al., 2011):

- Electrode locations were co-registered to the standard 1005 electrode template and warped to a 4-layer Boundary Element Method (BEM) model (Fuchs et al., 2002).
- For each IC topographical projection pattern, a single equivalent current dipole was autofit, optimising dipole location and orientation to minimise residual variance. For justification of this method, see Delorme et al. (2012).
- The effectiveness of dipole fitting was assessed through contrasting IC topography maps with dipole projections (residual variance) and estimating dipole location relative to the BEM template (Fuchs et al., 2002). In addition, a residual variance threshold of 20% was used to select for components with an acceptable dipole model fit, followed by filtering for remaining dipoles with valid locations. This resulted in total of 566/2852 ICs (19.8%) with less than 20% residual variance and equivalent dipole locations within the MNI brain mask that were included for source-based sensorimotor identification and k-means IC clustering.

### Spectral frequency analyses

The frequency decomposition of EEG data typically demonstrates a distinctive power distribution characterised by periodic narrowband oscillatory peaks combined with aperiodic 1/f-like activity (Donoghue et al., 2020; Gao et al., 2017; Gyurkovics et al., 2021). Therefore, periodic oscillations were estimated and separated from the aperiodic component using the Fieldtrip (version) (Oostenveld et al., 2011) implementation of Specparam (Donoghue et al., 2020), utilising code from Brainstorm (Tadel et al., 2011):

- Fast-Fourier transforms (FFT) were used to compute frequency representations for each electrode separately, creating uniform spectral resolution across frequencies. Spectra were estimated using 5 second, half-overlapping epochs with a Hanning-window taper (Delorme & Makeig, 2004; Furman et al., 2020; Lebrun, Ricci, et al., 2025), yielding a frequency resolution of 0.2 Hz. Spectra were estimated over 0.5 Hz to 100 Hz.
- Spectral frequency decompositions were averaged for each participant, session, time point, and electrode separately, and aperiodic estimation and subtraction were applied. Default spectral parameterisation parameters were used (Donoghue et al., 2020): peak detection threshold = 2.0 SD; peak widths = [0.5 12]; aperiodic mode = fixed; no maximum number of peaks; no minimum peak height.
- The quality of Specparam model fits was assessed using R^2^ and comparing the mean absolute fitting error between the fitted model and the observed PSD. This demonstrated excellent fits (R^2^ > 0.9) on average for sensor-level data when assessed globally and within each electrode cluster (Supplementary Table 1).
- The standard Specparam outputs of central frequency and peak amplitude for each periodic peak were quantified separately for each participant, session, time point, and electrode. These measurements were derived from the gaussian curves fitted to periodic data. For justification of this method, see Donoghue et al. (2020).
- PAF and α-band power were derived from the periodic peak with the highest amplitude and a central frequency falling within an 8 to 13 Hz range. When no peak was detected within this frequency band, an expanded range of 7 to 14 Hz was used (Haegens et al., 2014; Pérez-Elvira et al., 2026). This was necessary for 8 participants for sensor-level electrode grouping data. All global sensor-level PAF fell between the standard α-band range.

Global α-band oscillations and aperiodic components are displayed within Supplementary Figure 1a-c. Prior to removal of the aperiodic component, predominant posterior α-band oscillations were present at both parietal and occipital electrodes (Supplementary Figure 1a). However, following the isolation of periodic components, occipital α-band power was substantially reduced. This suggests that the higher magnitudes of α-band oscillations at occipital electrodes relative to anterior electrodes were dependent upon the aperiodic slope, which was indeed steeper towards the posterior (Supplementary Figure 1b).

### Spatial independence of PAF across sensor-level electrode groupings

Based upon previous research (Cavaleri et al., 2025; Cherin et al., 2026; Chowdhury et al., 2025; Furman et al., 2018; Furman et al., 2020, 2021; Furman et al., 2019; Lebrun, Ricci, et al., 2025; McLain et al., 2022), four regions of interest were defined for investigating α-band (8-13 Hz) periodic oscillations at the sensor-level alongside the global average across all electrodes: occipital (O1, Oz, O2); parietal (P1, Pz, P2); central (C1, Cz, C2); and frontal (F1, Fz, F2). However, given the likelihood of large volume-conduction biases when directly assessing sensor-level data, the following linear-mixed effects modelling was used to assess whether PAF genuinely differed spatially across electrode groupings prior to analyses of change in PAF over time or across repeated test sessions:

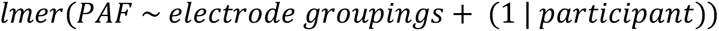

This revealed that there were statistically significant differences between electrode groupings (*F* (3, 1011) = 25.5, *p* < .001), although these explained only a very small proportion of the total variance (*marginal R^2^* = .02) compared to participant level random effects (*conditional R^2^* = .71), indicating that individual differences accounted for much of the observed variability in PAF.

Subsequently, pairwise comparisons between electrode groupings demonstrated a clear anterior-posterior distinction: frontal and central electrode groups showed similar PAF values (*Std. effect* = -.12 [−.3, .06], *p* = .55), as did parietal and occipital electrode groups (*Std. effect* = -.11 [−.29, .07], *p* = .57). However, all anterior-posterior comparisons (frontal-occipital; frontal-parietal; central-occipital; central-parietal) were statistically significant (*all Std. effect* > -.41 [−.06, -.23], *all p* < .001) indicating substantial differences.

In addition, separate linear models were fit independently for each electrode grouping to assess whether sensor-level data at these electrodes were primarily driven the same global variance:

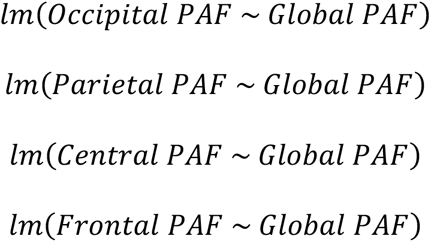

These models demonstrated that global oscillations significantly influenced the PAF of all electrode groupings (*all F* (1, 228) > 326.85, *all p* < .001) and captured a substantial, but not total, proportion of variance (59-84%) in each case (occipital electrodes: R^2^ = .84; parietal electrodes: R^2^ = .71; central electrodes: R^2^ = .79; frontal electrodes: R^2^ = .59). Therefore, whilst there was considerable shared variance between sensor-level electrode groupings and global PAF, substantial regional-specific variance remained.

These analyses were repeated for α-band power, demonstrating similar spatial variability across electrode groupings (*F* (3. 1011) = 55.08, *p* < .001) that only explained a small amount of variance (*marginal R^2^* = .03) relative to participant level random effects (*conditional R^2^* = .83). Furthermore, frontal and central electrode groupings demonstrated similarity (*Std. effect* = -.2 [−.38, -.02], *p* = .12), whilst anterior-posterior comparisons were all statistically significant (*all Std. effect* > -.52 [−.7, -.34], *all p* < .001). However, posterior and occipital electrodes were also significantly different (*Std. effect*= -.28 [−.46, -.1], *p* = .02), suggesting that parietal and occipital α-band power were distinct. In addition, the independent linear models used to assess the effect of global α-band power on electrode groupings confirmed a significant influence (*all F* (1, 228) > 724.38, *all p* < .001) and considerable shared variance (76-87%) in all cases (occipital electrodes: R^2^ = .87; parietal electrodes: R^2^ = .79; central electrodes: R^2^ = .76; frontal electrodes: R^2^ = .86), although substantial distinctions remained.

Together, these data demonstrate that when assessing α-band oscillations both region-specific variability and a substantial global signal shared across electrode groupings are present. Furthermore, two distinct anterior (frontal/central) and posterior (parietal/occipital) topographical groupings were evident. These are displayed within Supplementary Figure 2.

Therefore, subsequent sensor-level analyses examined PAF and α-band power globally and across residualised electrode groupings to investigate changes over time, between sessions, and in relation to MPS. Residualisation was accomplished through applying resid from the stats package to the above global variance models for each electrode grouping respectively. These data therefore represent the relative changes within each electrode group after accounting for the global α-band signal. For consistency, the same analyses applied to non-residualised occipital, parietal, central, and frontal electrode data are also presented.

### Identification of sensorimotor independent components

To investigate the reliability and specificity of component-based PAF changes following the induction of SHA, three separate clustering approaches were used: template matching (Chowdhury et al., 2025; Furman et al., 2018; Furman et al., 2020, 2021; Furman et al., 2019; Viola et al., 2009); optimised k-means clustering (Delorme & Makeig, 2004; Handiru et al., 2024; Miyakoshi et al., 2021; Rousseeuw, 1987; Thammasan & Miyakoshi, 2020); and dipole localisation to sensorimotor regions (Desikan et al., 2006; Oostenveld et al., 2011). These approaches were compared based upon agreement in component selection, consistency of dipole location, and, following the spectral frequency analyses described above, the reliability of cluster α-band power and PAF across sessions and relationships to SHA development. Spectral parameterisation was applied to each component separately, which demonstrated good fits (R^2^ > 0.86) on average (Supplementary Table 2). The identification of sensorimotor components was conducted as follows:

- Template matching was accomplished using the EEGLAB corrmap plugin (Viola et al., 2009) which involved the selection of a single component from each dataset with the highest spatial correlation to a designated template in the manner of Chowdhury et al. (2025) and Furman et al. (2018). Initially this was based upon a visually identified component (Supplementary Figure 3a). However, as demonstrated by Viola et al. (2009), the use of a single-subject template introduces individual-specific variability based upon anatomical discrepancies between participants. Therefore, a second correlation stage was applied using the average of all components initially identified as a more representative and stable template. Notably, components were not manually rejected based upon spectral power to avoid issues of circular inference.
- Optimal k-means clustering was applied to components with less than 20% residual variance and dipole locations within the BEM template (Fuchs et al., 2002). This resulted in a total of 566/2852 ICs (19.8%). Dipole moments, orientations, and component topographies were used as preclustering inputs and clustering was optimised using Silhouette and Calinski-Harabasz algorithms to maximise the ratio of unique subjects per cluster (Delorme & Makeig, 2004; Handiru et al., 2024; Miyakoshi et al., 2021; Rousseeuw, 1987; Thammasan & Miyakoshi, 2020). Notably, spectral power was not included as a metric to avoid issues of circular inference. Of the 7 clusters identified (Supplementary Figure 3b), cluster 4 demonstrated a topographical signature equivalent to the sensorimotor components previously reported (Chowdhury et al., 2025; Furman et al., 2018; Furman et al., 2020, 2021; Furman et al., 2019).
- Source localisation of ICs involved the registration of equivalent current dipole locations to the Desikan-Killinay atlas (Desikan et al., 2006; Oostenveld et al., 2011) based upon the BEM template (Fuchs et al., 2002). Components with less than 20% residual variance within the pre- and post-central gyri were considered sensorimotor. Components were not manually rejected based upon spectral power to avoid issues of circular inference.
- Upon the identification of multiple components within the same dataset, the component with the largest α-band peak was selected for further analyses (Chowdhury et al., 2025; Furman et al., 2018; Furman et al., 2020). This was necessary for both k-means clustering (15 datasets) and source localisation (28 datasets) methods.

A limitation of the present analyses is that only eyes-closed resting-state data were acquired. Therefore, whilst ICA blind source separation was applied, the use of a functional finger movement sensorimotor marker as demonstrated by Lebrun, Ricci, et al. (2025) was impossible. Despite this, the majority of previous research considering altered PAF has concentrated on either purely resting-state data (no concurrent task engagement) across participant cohorts (e.g., control vs chronic pain) or resting-state data altered by tonic/acute nociception within-participants (Ahmad & Barkana, 2025; Cavaleri et al., 2025; Cherin et al., 2026; Chowdhury et al., 2025; El Zghir et al., 2024; Fauchon et al., 2022; Furman et al., 2018; Furman et al., 2020; Furman et al., 2019; Liberati, 2025; McLain et al., 2022; Michalareas et al., 2016; Mussigmann et al., 2022; Nir et al., 2010; Ploner et al., 2017). Therefore, it remains unclear whether the isolation of a movement-related IC as performed by Lebrun, Ricci, et al. (2025) effectively captures the same phenomenon.

Furthermore, magnetoencephalographical research has demonstrated that the altered PAF present within neuropathic pain can be expressed within the default mode network (DMN), salience network, and various stages of ascending lemniscal thalamic transmission nodes (Fauchon et al., 2022). Therefore, it stands to reason that the engagement of motor control may interrupt or obscure these signals, particularly when considering that the DMN is anticorrelated with action mode network activation (Dosenbach et al., 2025), the ascending nociceptive system is strongly modulated by motor engagement (Darrow et al., 2025; Gordon et al., 2023; Umeda et al., 2019), and the salience network is guided by action goal-directives (Dosenbach et al., 2025; Seeley, 2019; Seeley et al., 2007). Therefore, the present analyses applying ICA to purely resting-state data remains in line with the majority of previous research that has successfully investigated the specificity of PAF alterations in relation to chronic or ongoing pain (Ahmad & Barkana, 2025; Cavaleri et al., 2025; Cherin et al., 2026; Chowdhury et al., 2025; El Zghir et al., 2024; Fauchon et al., 2022; Furman et al., 2018; Furman et al., 2020; Furman et al., 2019; Liberati, 2025; McLain et al., 2022; Michalareas et al., 2016; Mussigmann et al., 2022; Nir et al., 2010; Ploner et al., 2017).

### Statistical Analyses

All statistical analyses were conducted using R (version 4.3.1) (R Core Team, 2023).

### Intraclass correlation coefficients

Test-retest reliability was assessed through intraclass correlation coefficients (ICC 3,1), using a two-way mixed-effects model with absolute agreement for single rater measurements (Koo & Li, 2016) implemented in the psych package (Revelle, 2018). ICCs were interpreted as follows:

- Values less than 0.5 were indicative of poor reliability.
- Values between 0.5 and 0.75 were indicative of moderate reliability.
- Values between 0.75 and 0.9 were indicative of good reliability. Values greater than 0.9 were indicative of excellent reliability.

### Bland Altman limits of agreement

Agreement between sessions was assessed using Bland-Altman analyses (Bland & Altman, 1986; Chhapola et al., 2015; De Vet et al., 2006; Koo & Li, 2016), quantifying bias and limits of agreement (mean differences ±1.96 SD) relative to the observed range of values across sessions. This was implemented in the blandr package (Datta, 2017).

### Linear mixed-effects modelling

To investigate HFS-induced changes in resting-state EEG responses, statistical models were fit and interrogated for each EEG feature separately.

Data were analysed using linear mixed-effects models implemented in the lme4 package (Bates et al., 2015). Model assumptions were evaluated using the performance package (Lüdecke et al., 2021), including visual inspection of normality, linearity, collinearity, and homoscedasticity. Where violations were detected, optimal transformations were identified using the bestNormalize package (Peterson & Peterson, 2020). This resulted in log-plus-one transformations for MPS data and ordered normalisation transformations for PAF data.

Data (*Y*) were subsequently modelled with fixed effects for time (*β1*), session (*β2*), age (*β3*), sex (*β4*), and their interactions (*β5*). Critically, interactions were separated for sex and age to avoid overfitting four-way comparisons. Random intercepts for each individual (*u_i_*) were included to account for a within-individuals repeated-measures design:

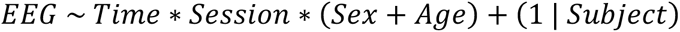

In addition, to investigate associations between resting-state EEG and MPS, EEG features were separately added as covariates to linear mixed-effect models as previously (Mugglestone, Ganis, et al., 2026) to assess their influence on reported pain intensity for each modality. Given the time-varying nature of resting-state EEG data immediately prior to pain rating blocks, it is critical to separate the between- and within-individuals variance so that valid inferences can be made (Hamaker & Muthén, 2020). Therefore, within each session, variance between-individuals was modelled as baseline values, whilst post-HFS changes relative to these baselines were considered within-individual values. No other predictors were included to minimise collinearity and focus interpretations:

For each model, conditional and marginal estimates of variance and effect size were calculated using mixed model R^2^ and partial Eta^2^ respectively. Fixed effects were evaluated via three-way within-individuals Type III ANOVA with Satterthwaite’s method for adjusting degrees of freedom implemented in the lmerTest package (Kuznetsova et al., 2015). Pairwise comparisons were conducted using estimated marginal means with Dunnett’s adjustment for multiple comparisons, implemented in the emmeans package (Lenth et al., 2025). Standardised effect sizes are also reported.

When applicable, Johnson–Neyman intervals were computed and visualised using the interactions package (Long & Long, 2021), applying a false discovery rate (FDR) at alpha = .05 to account for multiple comparisons (Esarey & Sumner, 2018). When necessary for visualisation, contrast-coding (−1; +1) was applied to the main predictor. However, this did not alter statistical significance or interpretation and was only implemented to facilitate visualisation.

## 5.2.4 Results

### Demographic characteristics

A total of 35 participants were recruited to the study, however, following an interim analysis of *n* = 8 participants, discrepancies with the accuracy of pinprick EEG registration was noticed leading to a halt in data collection. Following resolution of this issue, 6 participants failed to fully complete or return for both experimental sessions, 1 participant was excluded due to the disclosure of exclusion criteria, and 1 participant was excluded due to extreme artifacts within the EEG recording resulting in a final sample of 27 participants (female = 19).

### Resting-state EEG features

To demonstrate the consistency of EEG data according to prior demonstrations of resting-state EEG features, grand averaging across time points was conducted (Ahmad & Barkana, 2025; Cavaleri et al., 2025; Cherin et al., 2026; Chowdhury et al., 2025; El Zghir et al., 2024; Fauchon et al., 2022; Furman et al., 2018; Furman et al., 2020; Furman et al., 2019; Liberati, 2025; McLain et al., 2022; Michalareas et al., 2016; Mussigmann et al., 2022; Nir et al., 2010; Ploner et al., 2017). Therefore, Figure 1a-e displays a) MPS data demonstrating the induction and development of SHA as reported previously (Mugglestone, Ganis, et al., 2026), b) α-band peak power and PAF topographies, c) the global sensor-level α-band peak grand averaged across all time points, d) the relationship between change in global PAF post-HFS and MPS, e) component-level α-band peaks grand averaged across all time points for each component identification method separately.

**Figure 1.**
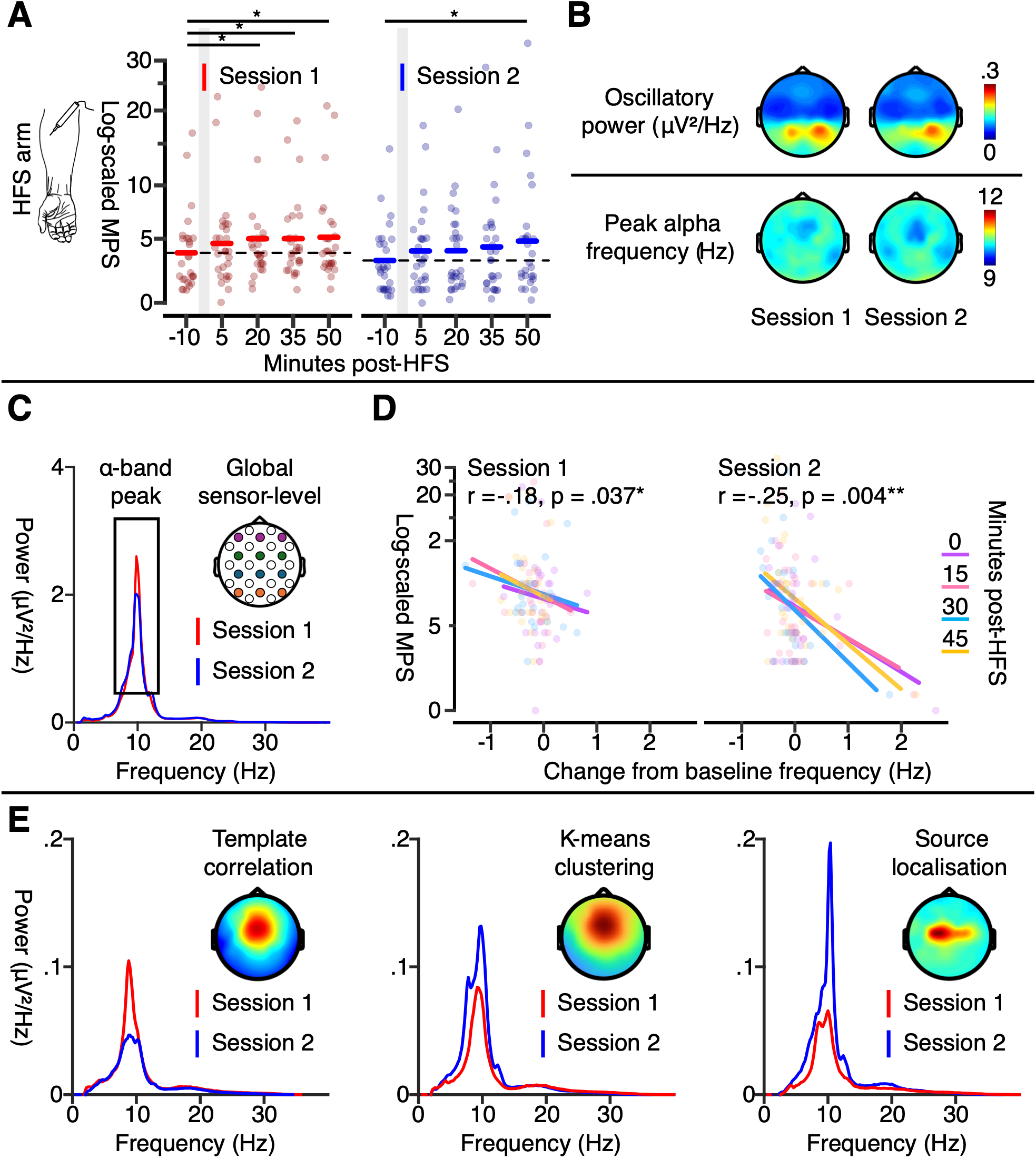
Primary α-band features. Raw data points are displayed as an average for each individual at each timepoint and session for the HFS arm MPS demonstrating the induction and development of SHA as previously reported. b) Topographies display α-band power (top) and PAF (bottom) grand averaged across time points for each session separately. c) The power spectrum plot displays the global sensor-level α-band peak grand averaged across time points for each session separately. d) Correlation plots display the relationships between change in global PAF post-HFS and MPS. e) Spectral frequency plots display the component-level α-band peaks grand averaged across time points for each session and component identification method separately. PAF = Peak alpha frequency; HFS = High-frequency stimulation.

Consistent with previous research, whilst α-band power and PAF varied across the topography (Figure 1b) a robust α-band peak was evident at the global average (Figure 1c) (Ahmad & Barkana, 2025; Cavaleri et al., 2025; Cherin et al., 2026; Chowdhury et al., 2025; El Zghir et al., 2024; Fauchon et al., 2022; Furman et al., 2018; Furman et al., 2020; Furman et al., 2019; Liberati, 2025; McLain et al., 2022; Michalareas et al., 2016; Mussigmann et al., 2022; Nir et al., 2010; Ploner et al., 2017). However, when compared across electrode groupings, substantial differences in amplitude and CF were present (Supplementary Figure 2). Previous investigations of PAF have evidenced some differences across electrode groupings within altered pain states (Cherin et al., 2026), however, the separation of global and region-specific variance was not considered. Therefore, the following analyses report both raw and residualised sensor-level data.

Furthermore, although conventional sensorimotor component topographies were demonstrated by all IC identification methods, substantial visual differences between sessions were evident for the averaged power spectrums (Figure 1e). In addition, there were large discrepancies in component selection that are reported further below.

### Test-retest reliability of sensor-level resting-state EEG

To investigate sensor-level EEG test-retest reliability at the global average and across electrode groupings, ICCs were calculated and are displayed within Table 1 alongside descriptive statistics concerning means, standard deviations, and ranges. Bland-Altman mean biases and LOA at baseline and throughout the HFS development time course are also reported within Table 1 and are displayed within Figure 2 alongside electrode grouping data. Notably, whilst all datasets demonstrated valid α-band peaks detectable by spectral parameterisation at the global level, this was not the case for all electrode groupings. Therefore, the sample sizes for these analyses were reduced (occipital: n = 24; parietal: n = 26; central; n = 25; frontal: n = 24).

**Figure 2.**
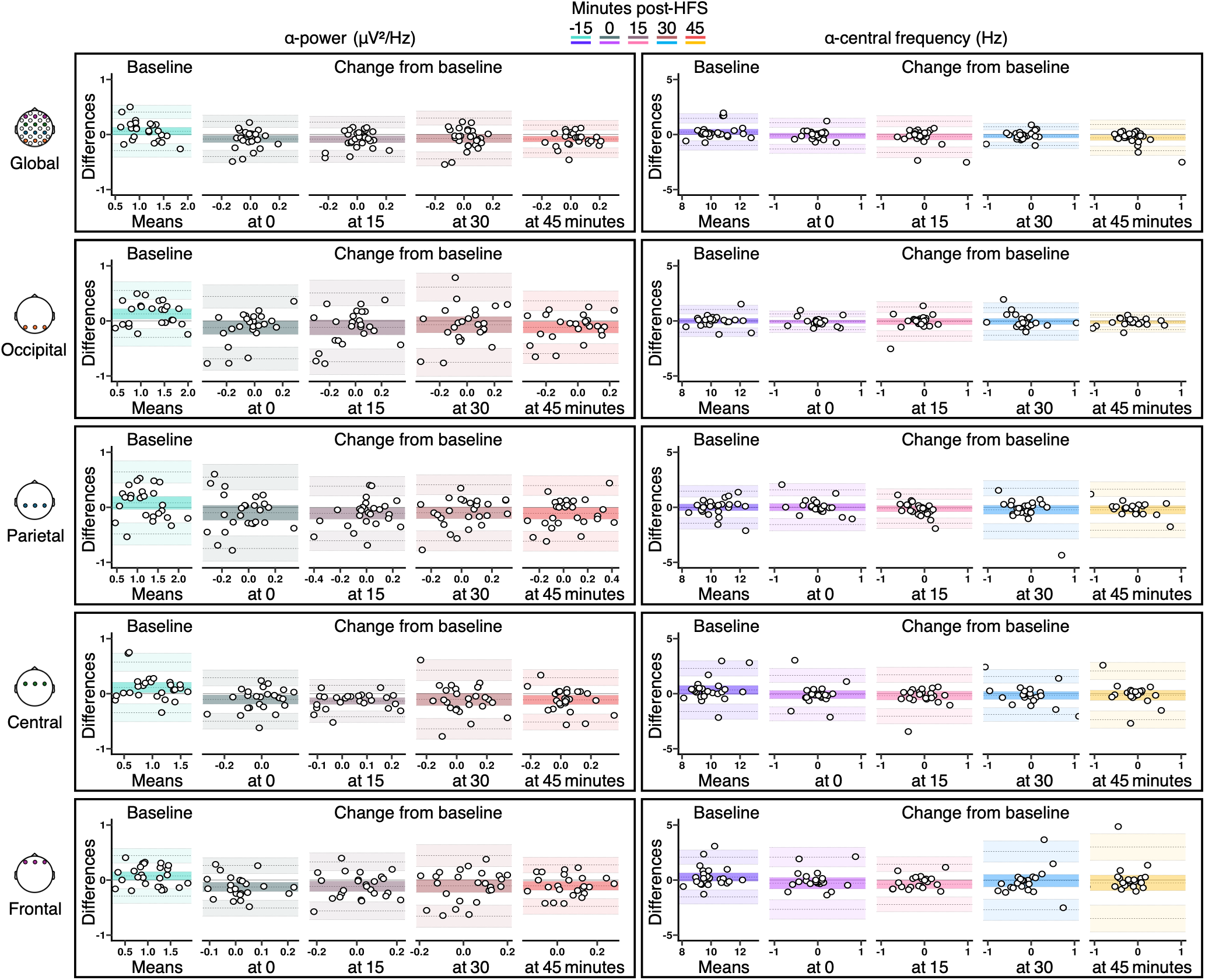
Sensor-level Bland-Altman α-band power and PAF assessments. At the global sensor-level and for each electrode grouping and α-band feature separately, Bland-Altman plots represent the mean bias (middle line), lower (bottom line), and upper (top line) limits of agreement, alongside 95% confidence intervals. Differences between measurements (y-axis) and the mean of paired measurements (x-axis) are displayed for each individual. HFS = High-frequency stimulation.

**Table 1.** Sensor-level α-band descriptive statistics and assessments of reliability and agreement. Mean values, standard deviations, and ranges were calculated separately for each session, time point, and sensor-level feature. Change from baseline scores were calculated as the absolute difference from baseline by subtracting the baseline values for each EEG response from their post-HFS counterparts. PAF = Peak alpha frequency; HFS = High-frequency stimulation.

| Key: | Value<br>(standard deviation) or [95% confidence interval] |  |  |  |  |
|---|---|---|---|---|---|
|  | **Raw data** | **Change from baseline (minutes)** |  |  |  |
| **Time post-HFS** | Baseline | 0 | 15 | 30 | 45 |
| **Power** (mV<sup>2</sup>/Hz) |  |  |  |  |  |
| **Global** |  |  |  |  |  |
| Session 1 mean | 1.1<br>(.4) | 0<br>(.1) | 0<br>(.1) | 0<br>(.1) | 0<br>(.1) |
| Session 1 range | .3 – 2 | -.4 – 0 | -.4 – 0 | -.4 – 0 | -.3 – 0 |
| Session 2 mean | 1<br>(.4) | 0<br>(.1) | 0<br>(.1) | .1<br>(.1) | .1<br>(.1) |
| Session 2 range | .4 – 2 | -.1 – 0 | -.1 – 0 | -.2 – 0 | -.1 – 0 |
| ICC | .89<br>[.78 .95] | 0<br>[-.37 .37] | .19<br>[-.2 .53] | 0<br>[-.37 .37] | .38<br>[. .66] |
| Bias | .06<br>[-.01 .13] | -.08<br>[-.14 -.02] | -.09<br>[-.15 -.02] | -.07<br>[-.15 0] | -.08<br>[-.13 -.03] |
| Upper | .41<br>[.29 .53] | .24<br>[.13 .35] | .22<br>[.12 .33] | .3<br>[.17 .43] | .17<br>[.08 .26] |
| Lower | -.29<br>[-.41 -.17] | -.4<br>[-.51 -.29] | -.4<br>[-.51 -.29] | -.44<br>[-.57 -.31] | -.33<br>[-.42 -.24] |
| **Occipital** |  |  |  |  |  |
| Session 1 mean | 1.3<br>(.5) | -.1<br>(.2) | -.1<br>(.3) | -0<br>(.3) | -.1<br>(.2) |
| Session 1 range | .4 – 2 | -.7 – 0 | -.7 – 0 | -.7 – 0 | -.6 – 0 |
| Session 2 mean | 1.1<br>(.5) | 0<br>(.1) | 0<br>(.2) | 0<br>(.2) | 0<br>(.2) |
| Session 2 range | .3 – 2 | -.3 – 0 | -.4 – 1 | -.5 – 0 | -.4 – 0 |
| ICC | .89<br>[.78 .95] | 0<br>[-.37 .37] | .05<br>[-.33 .42] | 0<br>[-.37 .37] | .27<br>[-.11 .59] |
| Bias | .13<br>[.03 .22] | -.12<br>[-.24 0] | -.12<br>[-.25 .02] | -.07<br>[-.22 .08] | -.12<br>[-.22 -.01] |
| Upper | .55<br>[.39 .71] | .44<br>[.23 .65] | .51<br>[.27 .74] | .61<br>[.36 .87] | .37<br>[.19 .54] |
| Lower | -.3<br>[-.46 -.14] | -.69<br>[-.9 -.47] | -.74<br>[-.98 -.51] | -.75<br>[-1 -.5] | -.6<br>[-.77 -.42] |
| **Parietal** |  |  |  |  |  |
| Session 1 mean | 1.2<br>(.4) | -.1<br>(.2) | -.1<br>(.2) | 0<br>(.2) | 0<br>(.3) |
| Session 1 range | .3 – 2 | -.6 – 0 | -.7 – 0 | -.6 – 0 | -.5 – 1 |
| Session 2 mean | 1.1<br>(.5) | 0<br>(.2) | .1<br>(.2) | .1<br>(.2) | .1<br>(.2) |
| Session 2 range | .5 – 2 | -.6 – 1 | -.2 – 0 | -.3 – 0 | -.3 – 1 |
| ICC | .79<br>[.6 .9] | 0<br>[-.37 .37] | .19<br>[-.2 .53] | .19<br>[-.2 .53] | .36<br>[-.2 .65] |
| Bias | .08<br>[-.04 .2] | -.10<br>[-.23 .03] | -.11<br>[-.21 -.01] | -.10<br>[-.21 0] | -.11<br>[-.21 -.01] |
| Upper | .64<br>[.44 .84] | .55<br>[.32 .78] | .38<br>[.21 .56] | .41<br>[.23 .59] | .39<br>[.21 .57] |
| Lower | -.48<br>[-.69 -.28] | -.75<br>[-.98 -.52] | -.61<br>[-.78 -.43] | -.61<br>[-.79 -.43] | -.61<br>[-.79 -.43] |
| **Central** |  |  |  |  |  |
| Session 1 mean | 1.1<br>(.4) | -.1<br>(.2) | 0<br>(.1) | 0<br>(.2) | 0<br>(.1) |
| Session 1 range | .3 – 2 | -.5 – 0 | -.3 – 0 | -.5 – 0 | -.2 – 0 |
| Session 2 mean | .9<br>(.4) | .0<br>(.1) | .1<br>(.1) | .1<br>(.2) | .1<br>(.2) |
| Session 2 range | .2 – 2 | -.2 – 0 | -.1 – 0 | -.5 – 1 | -.3 – 1 |
| ICC | .82<br>[.65 .91] | .03<br>[-.35 .4] | .4<br>[.03 .67] | 0<br>[-.37 .37] | .11<br>[-.28 .46] |
| Bias | .11<br>[.02 .21] | -.11<br>[-.19 -.02] | -.13<br>[-.19 -.07] | -.10<br>[-.21 0] | -.11<br>[-.19 -.03] |
| Upper | .57<br>[.41 .74] | .29<br>[.14 .43] | .15<br>[.05 .26] | .43<br>[.24 .62] | .29<br>[.15 .44] |
| Lower | -.34<br>[-.51 -.18] | -.5<br>[-.64 -.36] | -.42<br>[-.52 -.31] | -.64<br>[-.83 -.45] | -.51<br>[-.66 -.37] |
| **Frontal** |  |  |  |  |  |
| Session 1 mean | 1.1<br>(.4) | 0<br>(.1) | 0<br>(.2) | 0<br>(.2) | 0<br>(.1) |
| Session 1 range | .2 – 2 | -.3 – 0 | -.5 – 0 | -.4 – 0 | -.3 – 0 |
| Session 2 mean | 1<br>(.4) | .1<br>(.1) | .1<br>(.2) | .1<br>(.2) | .1<br>(.1) |
| Session 2 range | .3 – 2 | -.2 – 0 | -.3 – 0 | -.2 – 0 | -.2 – 0 |
| ICC | .9<br>[.8 .95] | 0<br>[-.37 .37] | .04<br>[-.34 .41] | 0<br>[-.37 .37] | .13<br>[-.26 .48] |
| Bias | .08<br>[0 .15] | -.12<br>[-.21 -.04] | -.11<br>[-.21 -.02] | -.11<br>[-.22 .01] | -.1<br>[-.18 -.02] |
| Upper | .44<br>[.31 .57] | .26<br>[.12 .41] | .33<br>[.17 .49] | .44<br>[.24 .64] | .27<br>[.13 .41] |
| Lower | -.29<br>[-.42 -.16] | -.51<br>[-.65 -.37] | -.56<br>[-.72 -.40] | -.66<br>[-.86 -.46] | -.48<br>[-.62 -.34] |
| **PAF** (Hz) |  |  |  |  |  |
| **Global** |  |  |  |  |  |
| Session 1 mean | 10.4<br>(1.2) | 0<br>(.3) | -.2<br>(.3) | -.2<br>(.4) | -.3<br>(.2) |
| Session 1 range | 8. – 13 | -.7 – 1 | -1.3 – 1 | -1.5 – 1 | -.8 – 0 |
| Session 2 mean | 10.2<br>(1) | .1<br>(.6) | 0<br>(.6) | -.1<br>(.5) | 0<br>(.6) |
| Session 2 range | 8.6 – 13 | -.5 – 3 | -.5 – 2 | -.6 – 2 | -.6 – 2 |
| ICC | .84<br>[.68 .92] | .13<br>[-.26 .48] | 0<br>[-.37 .37] | .49<br>[.14 .73] | 0<br>[-.37 .38] |
| Bias | .24<br>[-.01 .49] | -.12<br>[-.36 .12] | -.21<br>[-.49 .08] | -.14<br>[-.31 .03] | -.29<br>[-.53 -.05] |
| Upper | 1.47<br>[1.04 1.9] | 1.06<br>[.65 1.48] | 1.2<br>[.71 1.7] | .71<br>[.42 1.01] | .9<br>[.48 1.31] |
| Lower | -.99<br>[-1.42 -.56] | -1.3<br>[-1.71 -.89] | -1.62<br>[-2.11 -1.13] | -.99<br>[-1.28 -.69] | -1.48<br>[-1.9 -1.06] |
| **Occipital** |  |  |  |  |  |
| Session 1 mean | 10.4<br>(1.2) | 0<br>(.3) | -.1<br>(.5) | -.3<br>(.5) | -.2<br>(.4) |
| Session 1 range | 8 – 13 | -.9 – 1 | -2.1 – 1 | -1.9 – 1 | -1.4 – 1 |
| Session 2 mean | 10.3<br>(1.1) | 0<br>(.4) | -.1<br>(.3) | -.2<br>(.6) | -.1<br>(.4) |
| Session 2 range | 8.5 – 13 | -.9 – 1 | -1 – 0 | -1.6 – 1 | -.7 – 1 |
| ICC | .89<br>[.78 .95] | .45<br>[.09 .7] | 0<br>[-.37 .37] | .33<br>[-.05 .63] | .66<br>[.38 .83] |
| Bias | 0<br>[-.22 .22] | -.08<br>[-.23 .08] | -.06<br>[-.34 .23] | -.05<br>[-.32 .21] | -.12<br>[-.26 .03] |
| Upper | 1.03<br>[.65 1.42] | .64<br>[.37 .91] | 1.25<br>[.76 1.74] | 1.19<br>[.73 1.66] | .55<br>[.30 .8] |
| Lower | -1.03<br>[-1.42 -.65] | -.79<br>[-1.06 -.52] | -1.37<br>[-1.86 -.88] | -1.3<br>[-1.77 -.84] | -.78<br>[-1.03 -.53] |
| **Parietal** |  |  |  |  |  |
| Session 1 mean | 10.5<br>(1.1) | 0<br>(.6) | -.2<br>(.2) | -.2<br>(.7) | -.2<br>(.4) |
| Session 1 range | 8.2 – 13 | -2.3 – 1 | -.7 – 0 | -2 – 2 | -1.4 – 1 |
| Session 2 mean | 10.4<br>(1.1) | -.1<br>(1.1) | -.1<br>(.6) | -.1<br>(.8) | 0<br>(1) |
| Session 2 range | 8.7 – 13 | -3.6 – 3 | -2 – 1 | -1.8 – 3 | -2.4 – 3 |
| ICC | .77<br>[.55 .89] | .58<br>[.26 .78] | 0<br>[-.37 .37] | 0<br>[-.37 .37] | .22<br>[-.16 .55] |
| Bias | 0<br>[-.3 .3] | .01<br>[-.32 .34] | -.12<br>[-.4 .15] | -.22<br>[-.62 .18] | -.22<br>[-.61 .16] |
| Upper | 1.46<br>[.94 1.98] | 1.59<br>[1.03 2.16] | 1.21<br>[.73 1.68] | 1.73<br>[1.03 2.42] | 1.65<br>[.98 2.31] |
| Lower | -1.46<br>[-1.98 -.94] | -1.57<br>[-2.14 -1.01] | -1.45<br>[-1.93 -.98] | -2.17<br>[-2.87 -1.48] | -2.1<br>[-2.76 -1.43] |
| **Central** |  |  |  |  |  |
| Session 1 mean | 10<br>(1.3) | .1<br>(.7) | 0<br>(.7) | 0<br>(.6) | -.1<br>(.5) |
| Session 1 range | 7.9 – 14 | -1.4 – 2 | -2.1 – 2 | -.9 – 2 | -1.5 – 1 |
| Session 2 mean | 9.7<br>(1) | .2<br>(.7) | .2<br>(.9) | .2<br>(1) | .1<br>(1) |
| Session 2 range | 8.3 – 12 | -2 – 2 | -2.7 – 2 | -2.3 – 2 | -2.1 – 3 |
| ICC | .64<br>[.35 .82] | .18<br>[-.21 .52] | .3<br>[-.08 .61] | .41<br>[.04 .68] | .06<br>[-.32 .42] |
| Bias | .33<br>[-.07 .74] | -.08<br>[-.45 .28] | -.17<br>[-.57 .23] | -.16<br>[-.52 .2] | -.14<br>[-.6 .31] |
| Upper | 2.3<br>[1.6 2.99] | 1.66<br>[1.03 2.3] | 1.71<br>[1.02 2.4] | 1.57<br>[.96 2.19] | 2.06<br>[1.28 2.85] |
| Lower | -1.63<br>[-2.33 -.93] | -1.83<br>[-2.47 -1.19] | -2.05<br>[-2.74 -1.37] | -1.89<br>[-2.51 -1.27] | -2.35<br>[-3.13 -1.56] |
| **Frontal** |  |  |  |  |  |
| Session 1 mean | 10.2<br>(1.2) | 0<br>(.6) | -.2<br>(.6) | -.1<br>(.8) | -.2<br>(.7) |
| Session 1 range | 7.8 – 14 | -1.2 – 2 | -1.1 – 1 | -1.3 – 2 | -1.5 – 2 |
| Session 2 mean | 9.9<br>(1.3) | .1<br>(1.2) | .2<br>(1.1) | -.1<br>(1.3) | 0<br>(1.4) |
| Session 2 range | 8.3 – 14 | -2 – 4 | -2 – 4 | -2.7 – 3 | -2.9 – 4 |
| ICC | .75<br>[.51 .88] | .08<br>[-.31 .44] | .41<br>[.05 .68] | .18<br>[-.21 .52] | 0<br>[-.37 .37] |
| Bias | .28<br>[-.09 .65] | -.29<br>[-.8 .22] | -.38<br>[-.77 -0] | -.05<br>[-.61 .5] | -.25<br>[-.95 .45] |
| Upper | 2.09<br>[1.44 2.73] | 2.09<br>[1.2 2.98] | 1.44<br>[.77 2.11] | 2.6<br>[1.63 3.56] | 3<br>[1.79 4.21] |
| Lower | -1.53<br>[-2.18 -.89] | -2.66<br>[-3.55 -1.78] | -2.21<br>[-2.88 -1.54] | -2.7<br>[-3.67 -1.74] | -3.5<br>[-4.71 -2.29] |

At baseline, global power (ICC = .89) and PAF (ICC = .84) demonstrated good ICCs (Koo & Li, 2016). These high ICCs reflect the stability of individual differences across repeated test sessions. For PAF, this was accompanied by relatively small mean bias (.24 Hz) and limits of agreement (−.99 Hz to .44 Hz) relative to the assessed frequency band (7 Hz to 15 Hz). Alternatively, for power, whilst relatively small mean biases (.06 μV²/Hz) were present, there were larger limits of agreement (−.29 μV²/Hz to .41 μV²/Hz) relative to the range of observed data across sessions (.3 μV²/Hz to 2 μV²/Hz).

Furthermore, electrode groupings demonstrated good baseline reliability for power (occipital: ICC = .89; parietal: ICC = .79; central: ICC = .82; frontal: ICC = .9), and moderate-to-good reliability for PAF (occipital: ICC = .89; parietal: ICC = .77; central: ICC = .64; frontal: ICC = .75). When compared to global power, this was accompanied by relatively similar mean biases and LOA. Alternatively, PAF LOA were substantially wider for individual electrode groupings in comparison to the global average. However, whilst mean central and frontal electrode groupings demonstrated similar PAF biases to the global average, the mean bias occipital and parietal electrodes were almost zero, suggesting that there was almost no group-level shift in baseline PAF between sessions.

Therefore, overall, the strongest test-retest reliability of sensor-level resting-state EEG features were demonstrated at baseline for both power and PAF, with good reliability evident in almost every case. However, following HFS, change from baseline ICCs ranged from moderate to extremely poor, with a substantial number approaching zero (Table 1), indicating larger within-subject variability than between-subject variability. This was also accompanied by wider LOA, although mean biases remained relatively similar.

### Post-HFS changes in global sensor-level resting-state EEG across repeated test sessions

To investigate HFS-induced changes in global sensor-level resting-state EEG, mixed-effects modelling was used to assess change over time, between sessions, and the influences of age and sex for each EEG feature separately. These results are reported in Table 2 and displayed as group-level topographies and EEG responses in Figure 3 alongside electrode grouping data.

**Figure 3.**
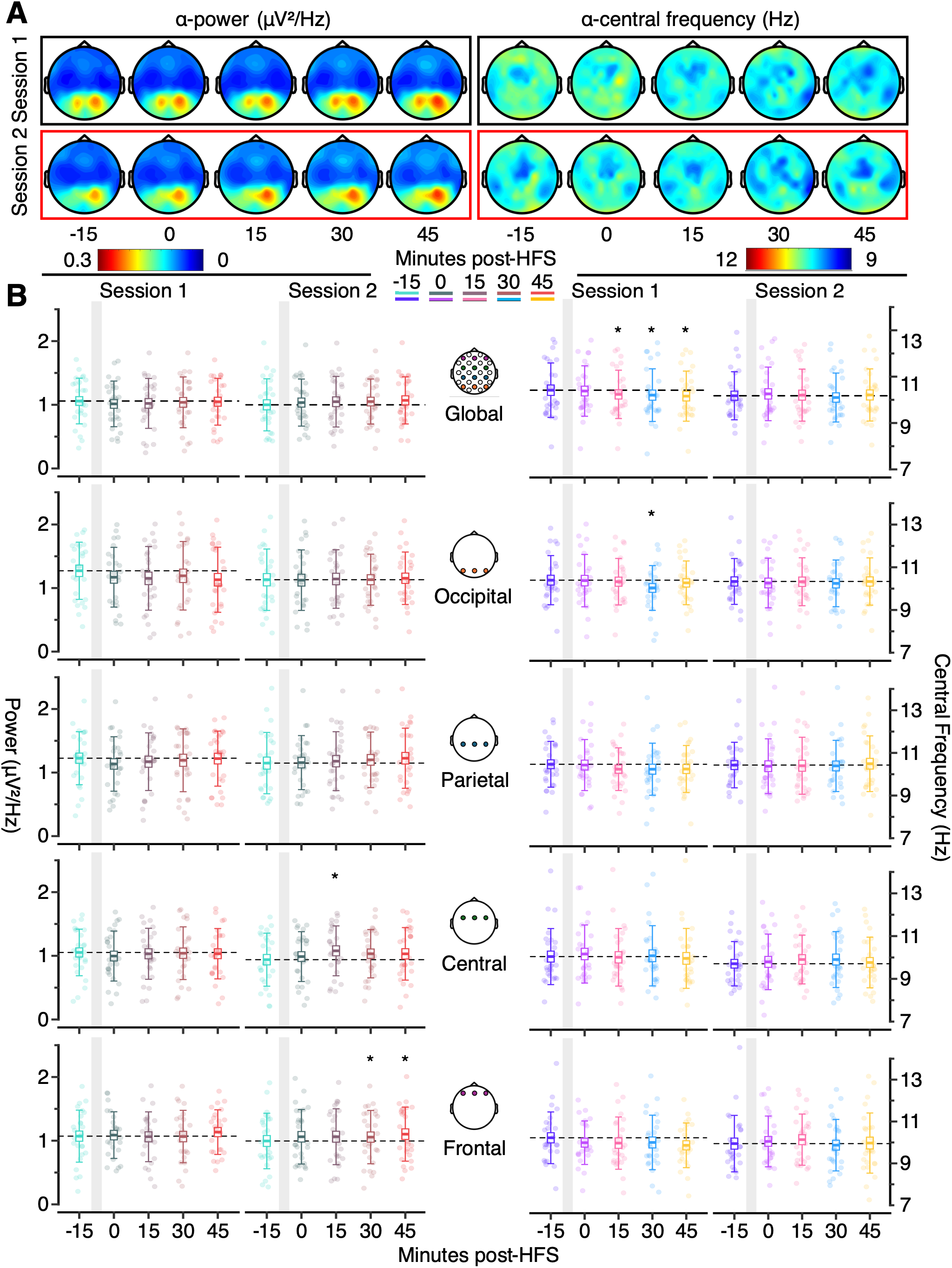
Sensor-level α -band power and PAF results. a) Topography plots represent the grand averaged sensor-level α-band power and PAF for each session respectively. b) Dot plots represent individual sensor-level α-band power and PAF data points alongside boxplots displaying group means, SEM, and 1 standard deviation. * indicates significance at alpha < .05. HFS = High-frequency stimulation.

**Table 2.** Global α-band mixed-effects modelling. The linear mixed effects modelling results for global α-band data are displayed for a) peak amplitude (power) b) PAF, and c-d) their relationships with MPS. PAF = Peak alpha frequency; HFS = High-frequency stimulation. N = 27, (female = 19).

| **a) Model assessment: Power**<br>Amplitude ~ time \* session \* (age + sex) + (1 \| subject) |  |  |  |
|---|---|---|---|
| Conditional R<sup>2</sup> = .95 | Marginal R<sup>2</sup> = .01 |  |  |
| **Fixed effects and interactions** |  |  |  |
| Predictor | *F* (DF) | *η2* [95% CI] | *p* |
| Time | F (4, 216) = .5 | <.01 [0, 1] | .74 |
| Session | F (1, 216) = .55 | <.01 [0, 1] | .46 |
| Age | F (1, 24) = .16 | <.01 [0, 1] | .69 |
| Sex | F (1, 24) = .04 | <.01 [0, 1] | .84 |
| Time: session | F (4, 216) = 1.84 | .03 [0, 1] | .12 |
| Time: age | F (4, 216) = .17 | <.01 [0, 1] | .95 |
| Time: sex | F (4, 216) = .65 | .01 [0, 1] | .63 |
| Session: age | F (1, 216) = .91 | <.01 [0, 1] | .34 |
| Session: sex | F (1, 216) = 6.91 | .03 [0, 1] | .01 |
| Time: session: age | F (4, 216) = 1.07 | .02 [0, 1] | .37 |
| Time: session: sex | F (4, 216) = .94 | .02 [0, 1] | .44 |
| **Between session comparisons: Session 2 – Session 1** |  |  |  |
| Minutes post-HFS | Estimate [95% CI] | Std. effect [95% CI] | *p* |
| -15 | -.07 [-.13, -.02] | -.79 [-1.39, -.2] | .01 |
| 0 | -.01 [-.06, .05] | -.06 [-.65, .53] | .84 |
| 15 | .02 [-.03, .08] | .24 [-.35, .83] | .42 |
| 30 | .01 [-.04, .07] | .13 [-.46, .71] | .68 |
| 45 | 0 [-.06, .05] | -.01 [-.6, .58] | .97 |
| **Session 1 comparisons to baseline: Post-HFS – Baseline** |  |  |  |
| Minutes post-HFS | Estimate [95% CI] | Std. effect [95% CI] | *p* |
| 0 | -.04 [-.11, .03] | -.41 [-1, .18] | .44 |
| 15 | -.05 [-.12, .02] | -.53 [-1.12, .06] | .24 |
| 30 | -.03 [-.1, .03] | -.37 [-.97, .22] | .52 |
| 45 | -.02 [-.08, .05] | -.17 [-.76, .41] | .9 |
| **Session 2 comparisons to baseline: Post-HFS – Baseline** |  |  |  |
| Minutes post-HFS | Estimate [95% CI] | Std. effect [95% CI] | *p* |
| 0 | .03 [-.04, .1] | .32 [-.27, .91] | .64 |
| 15 | .05 [-.02, .11] | .51 [-.08, 1.1] | .27 |
| 30 | .05 [-.02, .12] | .54 [-.05, 1.14] | .22 |
| 45 | .06 [-.01, .12] | .61 [.02, 1.2] | .14 |
| **b) Model assessment: PAF**<br>Frequency ~ time \* session \* (age + sex) + (1 \| subject) |  |  |  |
| Conditional R<sup>2</sup> = .89 | Marginal R<sup>2</sup> = .23 |  |  |
| **Fixed effects and interactions** |  |  |  |
| Predictor | *F* (DF) | *η2* [95% CI] | *p* |
| Time | F (4, 216) = 4.56 | .08 [.02, 1] | .001 |
| Session | F (1, 216) = 7.31 | .03 [0, 1] | .01 |
| Age | F (1, 24) = 0 | <.01 [0, 1] | .96 |
| Sex | F (1, 24) = 7.88 | .25 [.04, 1] | .01 |
| Time: session | F (4, 216) = 1.71 | .03 [0, 1] | .15 |
| Time: age | F (4, 216) = .1 | <.01 [0, 1] | .98 |
| Time: sex | F (4, 216) = 2.1 | .04 [0, 1] | .08 |
| Session: age | F (1, 216) = 3.29 | .01 [0, 1] | .07 |
| Session: sex | F (1, 216) = 3.18 | .01 [0, 1] | .08 |
| Time: session: age | F (4, 216) = 1.32 | .02 [0, 1] | .26 |
| Time: session: sex | F (4, 216) = .42 | <.01 [0, 1] | .79 |
| **Between session comparisons: Session 2 – Session 1** |  |  |  |
| Minutes post-HFS | Estimate [95% CI] | Std. effect [95% CI] | *p* |
| -15 | -.2 [-.33, -.06] | -.78 [-1.37, -.18] | .01 |
| 0 | -.18 [-.23, -.12] | -.66 [-1.25, -.07] | <.001 |
| 15 | -.05 [-.16, .06] | -.31 [-.9, .28] | .36 |
| 30 | -.05 [-.16, .06] | -.29 [-.88, .3] | .4 |
| 45 | .03 [-.03, .09] | .22 [-.37, .81] | .3 |
| **Session 1 comparisons to baseline: Post-HFS – Baseline** |  |  |  |
| Minutes post-HFS | Estimate [95% CI] | Std. effect [95% CI] | *p* |
| 0 | -.02 [-.24, .21] | -.08 [-.67, .51] | .99 |
| 15 | -.15 [-.28, -.02] | -.53 [-1.12, .06] | .02 |
| 30 | -.23 [-.44, -.02] | -.98 [-1.57, -.38] | .03 |
| 45 | -.23 [-.38, -.08] | -1.03 [-1.63, -.43] | .001 |
| **Session 2 comparisons to baseline: Post-HFS – Baseline** |  |  |  |
| Minutes post-HFS | Estimate [95% CI] | Std. effect [95% CI] | *P* |
| 0 | 0 [-.05, .05] | .04 [-.55, .63] | 1 |
| 15 | -.01 [-.06, .05] | -.06 [-.65, .53] | .99 |
| 30 | -.08 [-.19, .03] | -.49 [-1.08, .1] | .22 |
| 45 | 0 [-.07, .07] | -.03 [-.62, .56] | 1 |
| **c) Model assessment: Power influence on pain ratings**<br>Pain rating ~ time \* session \* (amplitude change + amplitude baseline) + (1 \| subject) |  |  |  |
| Conditional R<sup>2</sup> = .84 | Marginal R<sup>2</sup> = .09 |  |  |
| **Fixed effects and interactions** |  |  |  |
| Predictor | *F* (DF) | *η2* [95% CI] | *p* |
| Time | F (1, 238.16) = 28.58 | .11 [.05, 1] | <.001 |
| Session | F (1, 242.02) = 13.62 | .05 [.02, 1] | <.001 |
| Power change | F (1, 261.19) = 3.01 | .01 [0, 1] | .08 |
| Power baseline | F (1, 160.08) = 3.66 | .02 [0, 1] | .06 |
| Time: session | F (1, 238.23) = .3 | <.01 [0, 1] | .58 |
| Time: power change | F (1, 237.93) = .17 | <.01 [0, 1] | .68 |
| Time: power baseline | F (1, 237.83) = 2.53 | .01 [0, 1] | .11 |
| Session: power change | F (1, 24.15) = 6.17 | .03 [0, 1] | .01 |
| Session: power baseline | F (1, 242.2) = 1.64 | <.01 [0, 1] | .2 |
| Time: session: power change | F (1, 237.77) = .11 | <.01 [0, 1] | .74 |
| Time: session: power baseline | F (1, 238.06) = .08 | <.01 [0, 1] | .77 |
| **d) Model assessment: PAF influence on pain ratings**<br>Pain rating ~ time \* session \* (frequency change + frequency baseline) + (1 \| subject) |  |  |  |
| Conditional R<sup>2</sup> = .81 | Marginal R<sup>2</sup> = .09 |  |  |
| **Fixed effects and interactions** |  |  |  |
| Predictor | *F* (DF) | *η2* [95% CI] | *p* |
| Time | F (1, 243.3) = 27.67 | .1 [.05, 1] | <.001 |
| Session | F (1, 251.59) = 18.78 | .07 [.03, 1] | <.001 |
| Frequency change | F (1, 260.63) = 6.58 | .02 [0, 1] | .01 |
| Frequency baseline | F (1, 192.67) = .52 | <.01 [0, 1] | .47 |
| Time: session | F (1, 243.18) = .26 | <.01 [0, 1] | .61 |
| Time: frequency change | F (1, 243.2) = 0 | <.01 [0, 1] | .95 |
| Time: frequency baseline | F (1, 242.8) = 1.11 | <.01 [0, 1] | .29 |
| Session: frequency change | F (1, 245.35) = 9.08 | .04 [.01, 1] | .003 |
| Session: frequency baseline | F (1, 246.06) = 3.48 | .01 [0, 1] | .06 |
| Time: session: frequency change | F (1, 242.85) = .99 | <.01 [0, 1] | .32 |
| Time: session: frequency baseline | F (1, 242.46) = 1.38 | <.01 [0, 1] | .24 |

For global α-band power, no significant main effects were evident for either time (*F* (4, 216) = .5, *p* = .74) or session (*F* (1, 216) = .55, *p* = .46), although a significant pairwise comparison between sessions was demonstrated at baseline (*Std. effect* = -.79 [−1.39, - .2], *p* = .01), alongside a significant session-sex interaction (*F* (1, 216) = 6.91, *p* = .01). This suggests that at baseline, global α-band power substantially shifted between sessions for male participants (*F* (1, 180) = 5.68, *p* = .02), but not for female participants (*F* (1, 180) = .96, *p* = .33).

Alternatively, for global PAF, both a significant negative time effect (*F* (4, 216) = 4.56, *p* = .001) and a significant negative session effect (*F* (1, 216) = 7.31, *p* = .01) were present, alongside a significant difference between sexes (*F* (1, 24) = 7.88, *p* = .01). Pairwise comparisons revealed that this session effect was primarily driven by differences at baseline (*Std. effect* = -.78 [−1.37, -.18], *p* = .01) and immediately following HFS (Std. effect = -.66 [−1.25, -.07], *p* < .001). Furthermore, changes from baseline post-HFS were only statistically significant during session 1 at 15 (*Std. effect* = -.53 [1.12, .06], *p* = .02), 30 (*Std. effect* = -.98 [−1.57, -.38], *p* = .03), and 45 (*Std. effect* = −1.03 [−1.63, -.43], *p* = .001) minutes post-HFS. Although a negative trend was visually apparent during session 2, peaking at 30 minutes post-HFS (*Std. effect* = -.49 [−1.08, .1], *p* = .22), no pairwise comparisons reached statistical significance. This suggests that the slowing of global PAF occurred both between sessions and following HFS-induction, although during session two, the magnitude of change over time was substantially reduced. Furthermore, differences in PAF between male and female participants were evident, with male participants demonstrating slower PAF (9.51 Hz) on average compared to female participants (10.39 Hz).

### Post-HFS changes in sensor-level resting-state EEG at distinctive electrode groupings across repeated test sessions

For each electrode grouping, mixed-effects modelling was used to investigate the regional specificity of α-band changes between repeated test sessions and following HFS, alongside the influences of age and sex. Full spectral data are displayed in Supplementary Figure 4a-d. However, given the likelihood of volume-conduction biases blurring inference between electrode groupings, statistical modelling analyses were also applied to residualised data. To preserve the flow of text analyses are presented in Supplementary Tables. Together, these demonstrate various changes in α-band power and PAF between sessions, often mediated by individual differences. However, there was no support for any specific region driving the post-HFS changes in PAF that were evident at a global level.

When α-band power was assessed at occipital electrodes (Supplementary Table 1a), a significant negative main effect was evident for session (F (1, 204) = 5.13, p = .03) driven by a pairwise comparison at baseline (Std. effect = -.89 [−1.51, -.26], p = .01). This suggests that occipital electrodes captured the between-sessions difference in global α-band power. This was supported by residualised data (Supplementary Table 2a) demonstrating that the session effect remained significant after removal of global variance (F (1, 180) = 5.96, p = .02). However, for residualised data, effect sizes were smaller, and no significant pairwise comparisons were evident.

In contrast, no significant main effects were evident for α-band power when assessed at parietal (Supplementary Table 3a), central (Supplementary Table 5a), or frontal (Supplementary Table 7a) electrodes, despite substantial changes over time and between sessions driving various significant pairwise comparisons and interaction effects, none of which survived residualisation. Two cases are of note: Firstly, the residualisation of central electrode data (Supplementary Table 5a) revealed a significant interaction between session and age (F (1, 180) = 5.01, p = .03), demonstrating that only older participants (> 44 years) fluctuated significantly between sessions. Secondly, a significant interaction between time and session was evident for frontal electrodes (F (4, 202) = 2.59, p = .04), driven by the baseline pairwise comparison demonstrating a significant reduction in α-band power during session 2 (Std. effect = -.79 [−1.39, -.19], p = .01).

Together, these data suggest that α-band power varied inconsistently between repeated test sessions and across time, although these fluctuations were likely driven by global changes as residualised data demonstrated no significant main effects or pairwise comparisons.

Alternatively, when PAF was assessed, central electrodes (Supplementary Table 5b) demonstrated a significant difference between sessions (F (1, 203) = 6.05, p = .01), which was influenced by age (F (1, 203) = 8.57, p = .01), indicating that younger participants (< 33 years) changed significantly. However, only the main session effect (F (1, 180) = 5.26, p = .02) survived residualisation (Supplementary Table 6b). Similarly, an interaction between age and session was evident at parietal (F (1, 214) = 1.22, p = .01) and frontal (F (1, 202) = 5.25, p = .02) electrodes (Supplementary Table 3b and Supplementary Table 7b, respectively), although the effect only persisted through residualisation (Supplementary Table 4b) for the parietal grouping (F (1, 180) = 9.95, p = .01). This suggests that the oscillatory activity captured at parietal electrodes were driving the age-related differences in how PAF changed between sessions. Furthermore, a general main effect of age was evident when occipital electrodes were residualised (F (1, 200) = 8.67, p = .01).

Given the similarities between occipital and parietal electrode groupings (Supplementary Figure 2), this suggests that the oscillatory activity captured by posterior electrodes may be influenced most by age.

In addition, sex significantly influenced frontal (F (1, 24) = 4.45, p < .05), parietal (F (1, 240) = 7.32, p = .01), and occipital (F (1, 240) = 9.51, p = .001) PAF. Although this only survived residualisation for the parietal grouping (F (1, 200) = 4.83, p = .04), demonstrating significantly slower PAF for male participants (9.57 Hz) when compared with female participants (10.55 Hz). Therefore, parietal electrodes may also best capture sex-related differences in PAF

Together, these data suggest that posterior electrodes were most sensitive to demographical influences on PAF. However, whilst PAF may have fluctuated between sessions and at specific time points, no single electrode grouping was sensitive to the post-HFS changes in PAF that were evident at the global average.

### Relationships between global sensor-level resting-state EEG and MPS

To investigate the relationship between global sensor-level resting-state EEG and the development time course of SHA as assessed through MPS, mixed-effects modelling was used to characterise changes in MPS over time and between sessions as previously (Ganis, et al., 2026), with the addition of baseline and change from baseline EEG data included as covariates to capture between- and within-individuals variance respectively (Hamaker & Muthén, 2020). These results are reported in Table 2.

When global α-band power was assessed, neither baseline amplitude (*F* (1, 261) = 3.01, *p* = .08) nor change from baseline (*F* (1, 16) = 3.66, *p* = .06) were directly associated with MPS. However, a significant interaction between change in α-band power and session was evident (*F* (1, 24) = 6.17, *p* = .01), alongside significant main effects for time (*F* (1, 238) = 28.58, *p* < .001) and session (*F* (1, 242) = 13.62, *p* < .001) as previously reported (Mugglestone, Ganis, et al., 2026), which demonstrate the induction of SHA and variations in pain ratings across repeated test sessions respectively. Therefore, whilst some co-variations between MPS and changes in global α-band power were evident, this was not uniform across sessions, as evidenced by a strong effect during session 2 (*Std. trend* = −3.36 [−5.85, -.86], *p* = .009), but no statistically significant relationship between MPS and changes in global α-band power during session 1 (*Std. trend* = .42 [−1.67, 2.5], *p* = .69). This may have been due to differences between sessions in the development time course of SHA as previously reported (Mugglestone, Ganis, et al., 2026).

Alternatively, global PAF change from baseline was significantly associated with MPS (*F* (1, 260) = 6.58, *p* = .01), suggesting that intra-individual differences in PAF tracked with MPS. Whilst a significant interaction between session and PAF change was evident (*F* (1, 245) = 9.08, *p* = .003), further investigation revealed that the direction of the relationship between MPS and PAF change was consistent between sessions although the magnitude of this effect was stronger during session 2 than session 1 (Figure 1d). Therefore, despite a significant difference in strength, changes to global PAF post-HFS tracked the development time course of SHA in a consistent manner across sessions. Furthermore, no significant predictive effect of PAF at baseline was present (*F* (1, 193) = .52, *p* = .47), aligning with previous evidence demonstrating that baseline PAF is not a useful predictor of subsequently-induced SHA (Lebrun, Ricci, et al., 2025).

### Relationships between sensor-level resting-state EEG and MPS at distinctive electrode groupings

Mixed-effects modelling was also used to investigate the relationship between regional specific α-band EEG features and MPS. The same analyses were applied to residualised EEG data. For all analyses, significant main effects of time (*F* (1, 232) = 24.86, *p* < .001) and session (*F* (1, 207) = 3.02, *p* < .001) were apparent, evidencing the induction of SHA and differences in MPS between sessions as previously reported (Mugglestone, Ganis, et al., 2026). Therefore, for brevity, main effects for time and session are not reported separately for each case in the main text below.

When baseline α-band power was assessed, no significant main effect influence on MPS was evident for the occipital (Supplementary Table 1c) or parietal (Supplementary Table 3c) electrode groupings regardless of residualisation (Supplementary Table 2c, Supplementary Table 4c, respectively). A significant relationship between baseline α-band power and MPS was evident at frontal electrodes (*F* (1, 169) = 8.64, *p* < .001), but similarly did not survive residualisation, suggesting that the signal carrying this effect was not specific to the frontal grouping. In contrast, despite no main effects prior to residualisation, a significant relationship between MPS and baseline power was evident for residualised central electrodes (*F* (1, 227) = 4.71, *p* = .03), alongside an interaction with the change in MPS between sessions (*F* (1, 21) = 4,13, *p* = .04). This suggests that after accounting for global α-band variance, inter-individual differences in oscillatory activity at baseline specific to central electrodes were related to MPS, and that whilst the magnitude of this relationship differed between sessions, the direction was consistent. However, as no relationship between baseline α-band power and changes in MPS over time were evident, these data do not support a link between α-band power at a specific electrode grouping and the development of SHA. Furthermore, no significant relationships between change in α-band power and MPS were evident.

A significant relationship between baseline PAF and MPS (*F* (1, 252) = 5.81, *p* = .02) was evident for the central electrode grouping (Supplementary Table 5c), which may suggest that these inter-individual differences were reflective of general pain perception. However, no interaction with change in MPS over time was present. Alternatively, various significant interactions between baseline PAF and changes in MPS over time and between sessions were evident at occipital (Supplementary Table 1d) and parietal electrodes (Supplementary Table 3d). However, as no significant main effect was evident for either grouping, this demonstrates that the influence of baseline PAF fluctuated substantially rather than presenting a consistent indicator of changes in MPS. Therefore, these analyses demonstrate that whist central PAF at baseline may reflect inter-individual variations in pain perception, this was not sensitive to SHA development, whilst the relationships between MPS and baseline PAF at posterior electrode groupings were highly variable. Furthermore, no main effects or interactions were present when the influence of residualised baseline PAF was assessed, suggesting that these effects were not driven by these specific electrode groupings.

Alternatively, when changes in PAF were assessed at frontal electrodes (Supplementary Table 7d), a significant influence of change from baseline PAF on MPS was evident (*F* (1, 240) = 4.41, *p* = .04), as with the global average. However, residualised frontal data demonstrated no significant relationship. This suggests that whilst frontal electrodes carry the global signal, this was not in isolation. Alternatively, significant interactions between changes in PAF and changes in MPS were evident at occipital (Supplementary Table 1d), parietal (Supplementary Table 3d), and central (Supplementary Table 5d) electrodes, with the central (Supplementary Table 6d) and occipital (Supplementary Table 2d) effects surviving residualisation. This might indicate that the specific PAF changes captured by these electrodes were related to MPS in a manner that was not uniform but developed at later time points (see Supplementary Table 2d and Supplementary Table 6d for all estimates of marginal trends) and therefore, may reflect altered cortical dynamics at the later-phase of SHA development. However, as the present data only characterised resting-state EEG until 45 minutes post-HFS, further investigation of later time courses is required to verify this hypothesis.

Together, these data suggest that certain aspects of altered MPS following HFS can be linked to α-band power and PAF at particular electrode groupings. Specifically, the residualised baseline α-band power from central electrodes reflected general inter-individual differences in pain perception, and whilst no consistent effect of PAF on MPS survived residualisation, the late-phase development of a relationship between change in PAF and MPS was evident at occipital and central electrodes, although this requires further validation across a longer time course.

### Agreement of sensorimotor independent component identification methods

The identification of sensorimotor ICs through template matching (Chowdhury et al., 2025; Furman et al., 2018; Furman et al., 2020; Viola et al., 2009) involved the selection of a single component for each dataset with the highest spatial correlation to a designated template. Initially, this resulted in a mean correlation of .79 when fitting to a manually selected component. However, a second stage using the average of all components initially identified is necessary to accurately account for individual differences in head shape and electrode placement (Viola et al., 2009). This second stage improved component selection with the mean correlation rising to .85 as the result of the 3 datasets with the lowest correlation changing component selection. Therefore, there was a 93% carryover from the single-subject template to the group-average template. Secondly, optimal k-means clustering resulted in 7 independent clusters (Supplementary Figure 3b) with an average of 42 datasets and 78 components per cluster, alongside 24 outlier components (> 3 standard deviations from any cluster) originating from 21 different datasets. 4 datasets did not contribute any ICs to the sensorimotor cluster and 15 datasets contributed more than one. This may reflect a fundamental limitation of k-means clustering as consistently identifying a single component per dataset is implausible. Alternatively, this may reflect genuine individual differences in sensorimotor α-band oscillations, potentially improving signal-to-noise by omitting datasets with no relevant components. Accordingly, when the sensorimotor cluster was considered, there was an agreement of 61.1% (session 1 = 55.6%; session 2 = 66.7) between ICs identified through template matching and k-means clustering.

Thirdly, the source localisation of ICs through the registration of equivalent current dipoles to the Desikan-Killinay atlas (Desikan et al., 2006; Oostenveld et al., 2011) identified at least one component within every dataset and 28 datasets with multiple sensorimotor dipoles. However, the average agreement with template matching was only 31.5% (session 1 = 33.3%; session 2 = 29.6%), demonstrating that the majority of components selected through template matching did not localise within the pre- or post-central gyri, and therefore, may not be strictly considered sensorimotor in origin. Alternatively, the overlap between ICs identified through source localisation and those identified through k-means clustering was 50% (session 1 = 51.9%; session 2 = 48.1%).

Overall, k-means clustering demonstrated the highest agreement with both template matching and source localisation, albeit separately. This is unsurprising as it incorporates both topographical and dipole features (Delorme & Makeig, 2004). However, as components identified via template matching did not consistently localise to sensorimotor gyri, k-means clustering may impair the identification of relevant components by blurring the distinction between topographical similarity and anatomical localisation, which seem to reflect distinct phenomena.

### Test-retest reliability of component-level resting-state EEG

To investigate component-level EEG test-retest reliability, ICCs and Bland-Altman mean biases and LOA were calculated at baseline and throughout the HFS development time course. These are reported alongside descriptive statistics concerning means, standard deviations, and ranges within Table 3. Bland-Altman mean biases and LOA are displayed within Figure 4. Additionally, the proportion of datasets with no contributing components or identifiable α-band peak are presented in Supplementary Figure 5.

**Figure 4.**
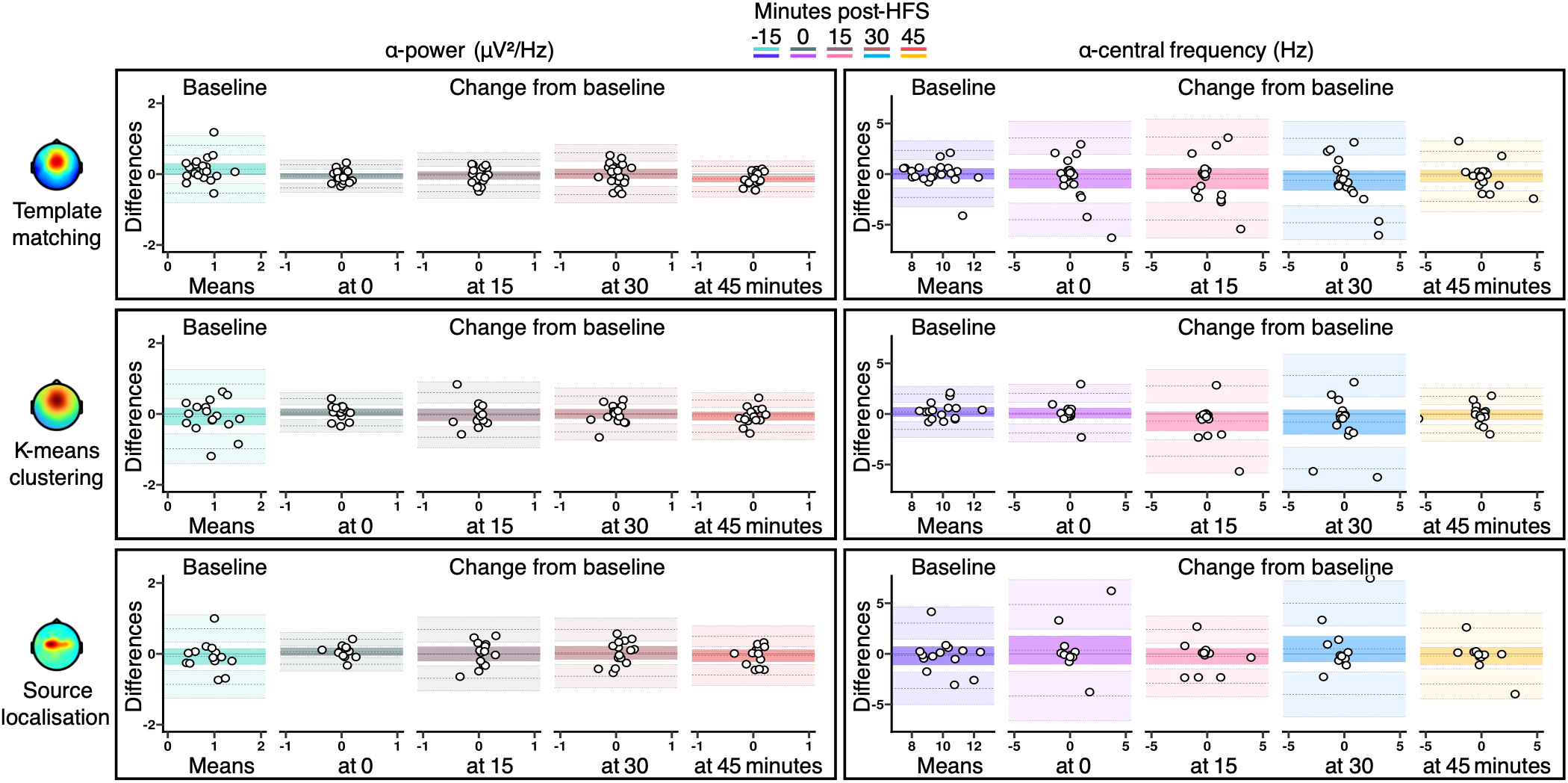
Component-level Bland-Altman assessments. For each method of sensorimotor component identification electrode grouping and α-band feature separately, Bland-Altman plots represent the mean bias (middle line), lower (bottom line), and upper (top line) limits of agreement, alongside 95% confidence intervals. Differences between measurements (y-axis) and the mean of paired measurements (x-axis) are displayed for each individual. PAF = Peak alpha frequency; HFS = High-frequency stimulation.

**Table 3.** Component-level α-band descriptive statistics and assessments of reliability. Mean values, standard deviations, and ranges were calculated separately for each session, time point, and component-level feature. Change from baseline scores were calculated as the absolute difference from baseline by subtracting the baseline values for each EEG response from their post-HFS counterparts. PAF = Peak alpha frequency; HFS = High-frequency stimulation. N = 27, (female = 19).

| Key: | Value<br>(standard deviation) or [95% confidence interval] |  |  |  |  |
|  | **Raw data** | **Change from baseline (minutes)** |  |  |  |
| **Time post-HFS** | Baseline | 0 | 15 | 30 | 45 |
|---|---|---|---|---|---|
| **Power** (mV<sup>2</sup>/Hz) |  |  |  |  |  |
| **Template matching** |  |  |  |  |  |
| Session 1 mean | .8<br>(.3) | 0<br>(.1) | 0<br>(.2) | 0<br>(.2) | 0<br>(.2) |
| Session 1 range | .3 - 2 | -.3 - 0 | -.2 - 0 | -.4 - 0 | -.4 - 0 |
| Session 2 mean | .7<br>(.3) | 0<br>(.1) | 0<br>(.1) | 0<br>(.2) | .1<br>(.1) |
| Session 2 range | .2 - 1 | -.2 - 0 | -.3 - 0 | -.4 - 0 | -.1 - 0 |
| ICC | .36<br>[-.02 .64] | .08<br>[-.3 .44] | 0<br>[-.37 .37] | 0<br>[-.37 .37] | .13<br>[-.26 .48] |
| Bias | .14<br>[-.01 .3] | -.06<br>[-.14 .02] | -.04<br>[-.15 .07] | .01<br>[-.13 .15] | -.14<br>[-.22 -.05] |
| Upper | .82<br>[.54 1.09] | .27<br>[.14 .4] | .41<br>[.22 .6] | .6<br>[.36 .84] | .23<br>[.08 .38] |
| Lower | -.53<br>[-.8 -.26] | -.39<br>[-.52 -.25] | -.49<br>[-.68 -.3] | -.58<br>[-.82 -.34] | -.5<br>[-.65 -.35] |
| **K-means clustering** |  |  |  |  |  |
| Session 1 mean | .8<br>(.4) | 0<br>(.1) | 0<br>(.2) | 0<br>(.2) | 0<br>(.2) |
| Session 1 range | .2 - 2 | -.3 - 0 | -.6 - 0 | -.6 - 0 | -.4 - 0 |
| Session 2 mean | .9<br>(.4) | 0<br>(.2) | 0<br>(.2) | 0<br>(.2) | 0<br>(.1) |
| Session 2 range | .2 - 2 | -.4 - 0 | -.8 - 0 | -.4 - 0 | -.2 - 0 |
| ICC | .41<br>[.04 .69] | .13<br>[-.26 .49] | .11<br>[-.28 .47] | .28<br>[-.11 .6] | .14<br>[-.25 .49] |
| Bias | -.07<br>[-.31 .17] | .05<br>[-.05 .15] | -.02<br>[-.2 .15] | 0<br>[-.13 .13] | -.06<br>[-.18 .06] |
| Upper | .84<br>[.43 1.26] | .43<br>[.26 .61] | .61<br>[.31 .91] | .51<br>[.27 .74] | .39<br>[.19 .6] |
| Lower | -.98<br>[-1.4 -.56] | -.34<br>[-.51 -.16] | -.66<br>[-.95 -.36] | -.51<br>[-.74 -.27] | -.51<br>[-.72 -.31] |
| **Source localisation** |  |  |  |  |  |
| Session 1 mean | .9<br>(.4) | .1<br>(.2) | 0<br>(.3) | .1<br>(.3) | .1<br>(.2) |
| Session 1 range | .3 - 2 | -.3 - 0 | -.7 - 1 | -.5 - 0 | -.3 - 0 |
| Session 2 mean | .9<br>(.4) | .1<br>(.2) | .1<br>(.2) | .1<br>(.2) | .1<br>(.2) |
| Session 2 range | .3 - 2 | -.4 - 1 | -.3 - 1 | -.3 - 1 | -.3 - 0 |
| ICC | .43<br>[.04 .7] | .46<br>[.09 .72] | 0<br>[-.39 .39] | 0<br>[-.39 .39] | .08<br>[-.32 .45] |
| Bias | -.07<br>[-.3 .15] | .06<br>[-.05 .17] | 0<br>[-.21 .2] | .03<br>[-.16 .21] | -.05<br>[-.22 .11] |
| Upper | .71<br>[.32 1.1] | .42<br>[.22 .61] | .68<br>[.33 1.04] | .69<br>[.36 1.01] | .49<br>[.2 .79] |
| Lower | -.86<br>[-1.25 -.47] | -.29<br>[-.49 -.1] | -.69<br>[-1.05 -.34] | -.63<br>[-.96 -.31] | -.6<br>[-.89 -.3] |
| **PAF** (Hz) |  |  |  |  |  |
| **Template matching** |  |  |  |  |  |
| Session 1 mean | 9.9<br>(1.6) | 0<br>(.8) | 0<br>(1.5) | -.4<br>(1.4) | -.1<br>(1.7) |
| Session 1 range | 7.8 - 13 | -1.1 - 2 | -3.2 - 4 | -5.6 - 2 | -5.9 - 3 |
| Session 2 mean | 9.5<br>(1.5) | .5<br>(1.8) | .3<br>(1.7) | .4<br>(2.1) | .2<br>(1.7) |
| Session 2 range | 7.2 - 13 | -2.4 - 7 | -2.4 - 6 | -2.6 - 6 | -3.7 - 6 |
| ICC | .72<br>[.47 .86] | 0<br>[-.37 .38] | .05<br>[-.33 .42] | .2<br>[-.19 .53] | .74<br>[.51 .87] |
| Bias | .03<br>[-.51 .57] | -.45<br>[-1.4 .49] | -.44<br>[-1.45 .57] | -.63<br>[-1.6 .34] | -.2<br>[-.79 .39] |
| Upper | 2.35<br>[1.41 3.29] | 3.59<br>[1.96 5.23] | 3.67<br>[1.91 5.43] | 3.54<br>[1.86 5.22] | 2.27<br>[1.24 3.29] |
| Lower | -2.3<br>[-3.24 -1.36] | -4.5<br>[-6.14 -2.87] | -4.55<br>[-6.31 -2.8] | -4.8<br>[-6.48 -3.12] | -2.67<br>[-3.7 -1.65] |
| **K-means clustering** |  |  |  |  |  |
| Session 1 mean | 1.4<br>(1.4) | 0<br>(.7) | -.2<br>(1.1) | -.3<br>(1.7) | -.4<br>(1.4) |
| Session 1 range | 8.6 - 13 | -1.1 - 2 | -3.2 - 2 | -5.6 - 3 | -5.9 - 2 |
| Session 2 mean | 9.8<br>(1.5) | .2<br>(1.7) | .1<br>(1.8) | .5<br>(1.9) | 0<br>(1.9) |
| Session 2 range | 7.8 - 14 | -2.1 - 7 | -5.1 - 6 | -2.1 - 6 | -5.4 - 6 |
| ICC | .81<br>[.62 .91] | .77<br>[.55 .89] | 0<br>[-.38 .38] | 0<br>[-.38 .38] | .87<br>[.73 .94] |
| Bias | .2<br>[-.25 .66] | .08<br>[-.43 .59] | -.72<br>[-1.66 .23] | -.8<br>[-2.01 .41] | -.07<br>[-.54 .4] |
| Upper | 1.94<br>[1.15 2.74] | 2.04<br>[1.14 2.93] | 2.75<br>[1.11 4.4] | 3.81<br>[1.7 5.91] | 1.73<br>[.91 2.55] |
| Lower | -1.53<br>[-2.33 -.74] | -1.88<br>[-2.77 -.98] | -4.19<br>[-5.84 -2.54] | -5.41<br>[-7.52 -3.3] | -1.86<br>[-2.68 -1.04] |
| **Source localisation** |  |  |  |  |  |
| Session 1 mean | 1.3<br>(1.4) | .4<br>(1.8) | -.2<br>(1.4) | .1<br>(1.8) | -.2<br>(.8) |
| Session 1 range | 8.1 - 13 | -.8 - 7 | -3.2 - 4 | -3. - 6 | -2.1 - 2 |
| Session 2 mean | 1.5<br>(2.0) | -.5<br>(2.0) | -.4<br>(1.9) | -.6<br>(1.2) | -.3<br>(2.0) |
| Session 2 range | 7.2 - 14 | -7.3 - 4 | -4.9 - 4 | -3.7 - 1 | -4. - 5 |
| ICC | .56<br>.22 .78] | .18<br>[-.22 .53] | .69<br>[.41 .85] | 0<br>[-.39 .39] | .59<br>[.26 .8] |
| Bias | -.19<br>[-1.1 .72] | .37<br>[-1.03 1.76] | -.25<br>[-1.02 .53] | .47<br>[-.8 1.74] | -.22<br>[-1.06 .63] |
| Upper | 3.04<br>[1.44 4.64] | 4.88<br>[2.44 7.32] | 2.4<br>[1.03 3.76] | 4.97<br>[2.75 7.2] | 2.53<br>[1.04 4.02] |
| Lower | -3.42<br>[-5.02 -1.83] | -4.15<br>[-6.59 -1.71] | -2.89<br>[-4.25 -1.52] | -4.03<br>[-6.25 -1.81] | -2.96<br>[-4.45 -1.48] |

At baseline, the α-band power of ICs identified through template correlation demonstrated poor reliability (ICC = .36) with wide LOA (−.53 μV²/Hz to .82 μV²/Hz) relative to the range of observed data across sessions (.3 μV²/Hz to 2 μV²/Hz). Alternatively, whilst PAF reliability was moderate (ICC = .72), LOA were extremely wide (−2.3 Hz to 2.35 HZ) relative to the typical α-band (8 Hz to 12 Hz). Furthermore, although a component was successfully identified for each dataset, 12 datasets demonstrated no valid α-band peak detectable by spectral parameterisation and therefore could not be included within these analyses.

K-means clustering demonstrated poor reliability for power (ICC = .41) but good reliability for PAF (ICC = .81), suggesting relatively more stable individual differences across sessions. However, this was accompanied by larger LOA for α-band power (−.98 μV²/Hz to .42 μV²/Hz), and although PAF LOA were slightly narrower (−1.53 Hz to 1.94 Hz) they remained extremely wide. Furthermore, 6 datasets contributed no components to the sensorimotor cluster and therefore could not be included within these analyses.

For ICs identified through source localisation, test-retest reliability was moderate to poor for both power (ICC = .43) and PAF (ICC = .56). These were accompanied by wide LOA for both power (−.86 μV²/Hz to .71 μV²/Hz) and PAF (−3.42 Hz to 3.04 Hz). Additionally, whilst source localisation identified at least one sensorimotor component for all datasets, no valid α-band peak detectable by spectral parameterisation was present for any sensorimotor components within 8 datasets.

Overall, at baseline, the test-retest reliability of α-band power was poor for all three methods, with wide LOA. In addition, LOA for PAF data were extremely wide, despite comparatively higher ICCs.

Following HFS, change from baseline ICCs ranged from moderate to extremely poor in most cases, accompanied by relatively wider LOA throughout. However, PAF reliability returned to near pre-HFS scores at 45 minutes in all cases, suggesting that component-level PAF may be more reliable for identifying late-phase changes induced by HFS.

For α-band power change from baseline, template matching ICCs in ascending time order were .08, 0, 0, and .13; K-means clustering ICCs were .13, .11, .28, and .14; source localisation ICCs were .46, 0, 0, and .08.

For PAF change from baseline, template matching ICCs in ascending time order were 0, .05, .2, and .74; K-means clustering ICCs were .77, 0, 0, and .87; Source localisation ICCs were .18, .69, 0, and .59.

Overall, the strongest test-retest reliability of component-level α-band power occurred at baseline, with substantially reduced ICCs when changes from baseline were assessed in most cases. However, whilst PAF ICCs dipped during the early temporal dynamics of SHA development, at 45 minutes post-HFS a return to baseline-level reliability was evident.

Therefore, component-level PAF methods may be best suited to characterising the late-phase expression of SHA. Furthermore, several participants demonstrated no identifiable α-band peak despite valid component selections through template matching and source localisation. In addition, many of these participants failed to contribute to the k-means sensorimotor cluster. This lack of consistent PAF identification across participants and sessions may reflect a fundamental weakness of these methods. However, as this concern was present across methodologies, the exclusion of these participants from the k-means cluster may instead demonstrate a sensitivity for selecting only valid sensorimotor α-band activity. Therefore, further analyses below assess the relationships between component-level resting-state EEG and MPS to clarify the accuracy of these methods for capturing changes in reported pain ratings throughout the HFS-induced SHA development time course.

### Post-HFS changes in component-level resting-state EEG across repeated test sessions

For each sensorimotor component identification method, mixed-effects modelling was used to investigate α-band and PAF changes between repeated test sessions and following HFS, alongside the influences of age and sex. These results are displayed within Figure 5. Together, these data demonstrate various changes in α-band power and PAF between sessions, with large amplitude related differences between sessions (Figure 1e) often mediated by individual differences. However, there was no support for any specific component identification method successfully capturing the post-HFS changes in PAF that were evident at the sensor-level. Furthermore, given the inconsistent selection of components and lack of identifiable α-band peaks for a number of datasets, only participants with data in both sessions were included (Supplementary Figure 5).

**Figure 5.**
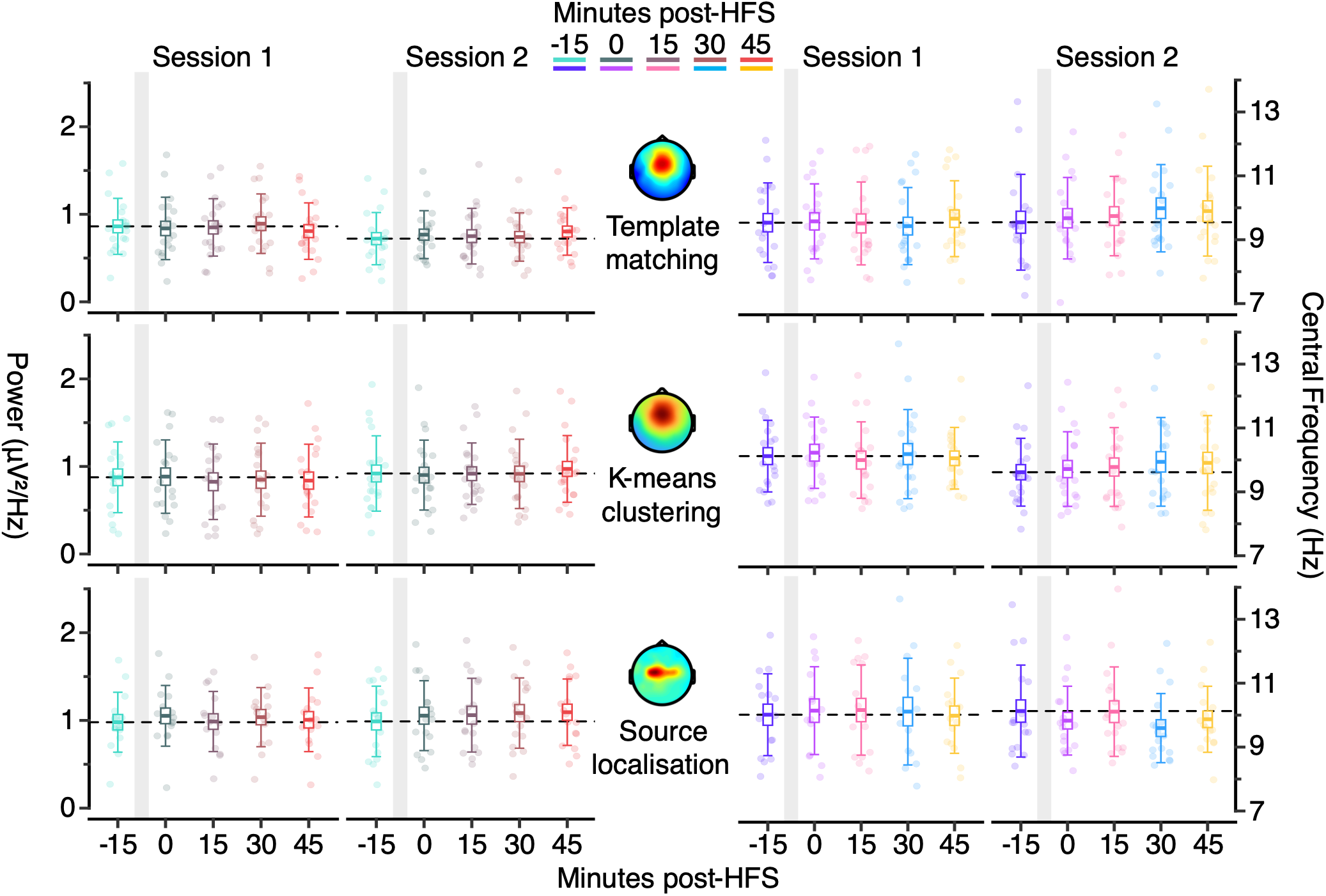
Component-level α-band power and PAF results. Dot plots represent individual sensor-level α-band power (top) and PAF (bottom) data points alongside boxplots displaying group means, SEM, and 1 standard deviation. No significant pairwise comparisons to baseline were evident. PAF = Peak alpha frequency; HFS = High-frequency stimulation.

For components identified through template matching (Table 4), a significant negative main effect was evident changes in α-band power between sessions (*F* (1, 144) = 7.31, *p* = .01), which was driven by significant pairwise comparisons at baseline (*Std. effect* = - .75 [−1.43, -.08], *p* <.001), and at 15 minutes (*Std. effect* = -.51 [−1.18, .16], *p* = .01) and 30 minutes (*Std. effect* = -.74 [−1.41, -.07], *p* <.001) post-HFS. This was mediated by age (*F* (1, 144) = 6.55, *p* = .01) as younger participants (< 25) experienced smaller fluctuations between sessions. However, no other main effects or interactions were significant. For template matching PAF, a significant main effect of session was again evident (*F* (1, 144) = 5.72, *p* = .02). However, no pairwise comparisons, further main effects, nor interactions were significant.

**Table 4.** Template matching α-band mixed-effects modelling. The linear mixed effects modelling results for template matching α-band data are displayed for a) peak amplitude (power) b) PAF, and c-d) their relationships with MPS. = High-frequency stimulation. N = 27, (female = 19).

| **a) Model assessment: Power**<br>Amplitude ~ time \* session \* (age + sex) + (1 \| subject) |  |  |  |
|---|---|---|---|
| Conditional R<sup>2</sup> = .64 | Marginal R<sup>2</sup> = .08 |  |  |
| **Fixed effects and interactions** |  |  |  |
| Predictor | *F* (DF) | *η2* [95% CI] | *p* |
| Time | F (4, 144) = .02 | <.01 [0, 1] | 1 |
| Session | F (1, 144) = 7.31 | .05 [.01, 1] | .01 |
| Age | F (1, 16) = .25 | .02 [0, 1] | .63 |
| Sex | F (1, 16) = .81 | .05 [0, 1] | .38 |
| Time: session | F (4, 144) = 1.12 | .03 [0, 1] | .35 |
| Time: age | F (4, 144) = .27 | <.01 [0, 1] | .9 |
| Time: sex | F (4, 144) = .19 | <.01 [0, 1] | .94 |
| Session: age | F (1, 144) = 6.55 | .04 [.01, 1] | .01 |
| Session: sex | F (1, 144) = .14 | <.01 [0, 1] | .7 |
| Time: session: age | F (4, 144) = .37 | .01 [0, 1] | .83 |
| Time: session: sex | F (4, 144) = .3 | <.01 [0, 1] | .88 |
| **Between session comparisons: Session 2 – Session 1** |  |  |  |
| Minutes post-HFS | Estimate [95% CI] | Std. effect [95% CI] | *p* |
| -15 | -.16 [-.24, -.07] | -.75 [-1.43, -.08] | <.001 |
| 0 | -.08 [-.27, .1] | -.36 [-1.03, .31] | .36 |
| 15 | -.13 [-.22, -.03] | -.51 [-1.18, .16] | .01 |
| 30 | -.16 [-.24, -.08] | -.74 [-1.41, -.07] | <.001 |
| 45 | .01 [-.2, .23] | .12 [-.55, .78] | .89 |
| **Session 1 comparisons to baseline: Post-HFS – Baseline** |  |  |  |
| Minutes post-HFS | Estimate [95% CI] | Std. effect [95% CI] | *p* |
| 0 | -.03 [-.25, .2] | -.14 [-.81, .53] | .98 |
| 15 | -.01 [-.13, .1] | -.1 [-.76, .57] | .98 |
| 30 | .01 [-.09, .12] | .07 [-.6, .73] | .98 |
| 45 | -.1 [-.22, .01] | -.41 [-1.08, .25] | .08 |
| **Session 2 comparisons to baseline: Post-HFS – Baseline** |  |  |  |
| Minutes post-HFS | Estimate [95% CI] | Std. effect [95% CI] | *p* |
| 0 | .05 [-.03, .12] | .25 [-.41, .92] | .38 |
| 15 | .01 [-.09, .12] | .15 [-.51, .82] | .98 |
| 30 | .01 [-.1, .11] | .08 [-.58, .75] | .99 |
| 45 | .07 [-.2, .33] | .46 [-.21, 1.13] | .88 |
| **b) Model assessment: PAF**<br>Frequency ~ time \* session \* (age + sex) + (1 \| subject) |  |  |  |
| Conditional R<sup>2</sup> = .47 | Marginal R<sup>2</sup> = .12 |  |  |
| **Fixed effects and interactions** |  |  |  |
| Predictor | *F* (DF) | *η2* [95% CI] | *p* |
| Time | F (4, 144) = .22 | <.01 [0, 1] | .93 |
| Session | F (1, 144) = 5.72 | .04 [0, 1] | .02 |
| Age | F (1, 16) = 1.21 | .07 [0, 1] | .29 |
| Sex | F (1, 16) = 1.54 | .09 [0, 1] | .23 |
| Time: session | F (4, 144) = .83 | .02 [0, 1] | .51 |
| Time: age | F (4, 144) = .57 | .02 [0, 1] | .69 |
| Time: sex | F (4, 144) = .87 | .02 [0, 1] | .48 |
| Session: age | F (1, 144) = 1.57 | .01 [0, 1] | .21 |
| Session: sex | F (1, 144) = .84 | <.01 [0, 1] | .36 |
| Time: session: age | F (4, 144) = 1.11 | .03 [0, 1] | .35 |
| Time: session: sex | F (4, 144) = .3 | <.01 [0, 1] | .88 |
| **Between session comparisons: Session 2 – Session 1** |  |  |  |
| Minutes post-HFS | Estimate [95% CI] | Std. effect [95% CI] | *p* |
| -15 | .03 [-.59, .66] | .06 [-.61, .73] | .91 |
| 0 | .32 [-.41, 1.05] | .37 [-.3, 1.03] | .38 |
| 15 | .47 [-.08, 1.02] | .54 [-.13, 1.21] | .09 |
| 30 | .79 [-2.11, 3.68] | .76 [.09, 1.43] | .59 |
| 45 | .13 [-.2, .45] | .18 [-.48, .85] | .44 |
| **Session 1 comparisons to baseline: Post-HFS – Baseline** |  |  |  |
| Minutes post-HFS | Estimate [95% CI] | Std. effect [95% CI] | *p* |
| 0 | 0 [-.65, .64] | -.01 [-.67, .66] | 1 |
| 15 | -.14 [-.88, .6] | -.16 [-.83, .5] | .94 |
| 30 | -.2 [-.74, .34] | -.21 [-.88, .45] | .73 |
| 45 | .15 [-.37, .66] | .13 [-.54, .8] | .84 |
| **Session 2 comparisons to baseline: Post-HFS – Baseline** |  |  |  |
| Minutes post-HFS | Estimate [95% CI] | Std. effect [95% CI] | *P* |
| 0 | .28 [-.71, 1.27] | .3 [-.37, .97] | .84 |
| 15 | .3 [-.46, 1.06] | .32 [-.35, .99] | .69 |
| 30 | .56 [-2.99, 4.1] | .49 [-.18, 1.15] | .96 |
| 45 | .24 [-.52, 1] | .26 [-.41, .92] | .8 |
| **c) Model assessment: Power influence on pain ratings**<br>Pain rating ~ time \* session \* (amplitude change + amplitude baseline) + (1 \| subject) |  |  |  |
| Conditional R<sup>2</sup> = .85 | Marginal R<sup>2</sup> = .08 |  |  |
| **Fixed effects and interactions** |  |  |  |
| Predictor | *F* (DF) | *η2* [95% CI] | *p* |
| Time | F (1, 17.76) = 14.91 | .08 [.03, 1] | <.001 |
| Session | F (1, 172.26) = 27.17 | .14 [.07, 1] | <.001 |
| Power change | F (1, 172.77) = 7.43 | .04 [.01, 1] | .01 |
| Power baseline | F (1, 182.55) = 6.55 | .03 [0, 1] | .01 |
| Time: session | F (1, 17.76) = .68 | <.01 [0, 1] | .41 |
| Time: power change | F (1, 171.07) = 1.83 | .01 [0, 1] | .18 |
| Time: power baseline | F (1, 17.8) = 1.13 | <.01 [0, 1] | .29 |
| Session: power change | F (1, 173.45) = .3 | <.01 [0, 1] | .59 |
| Session: power baseline | F (1, 173.58) = 2.62 | .01 [0, 1] | .11 |
| Time: session: power change | F (1, 171.01) = 1.15 | <.01 [0, 1] | .29 |
| Time: session: power baseline | F (1, 17.8) = .66 | <.01 [0, 1] | .42 |
| **d) Model assessment: PAF influence on pain ratings**<br>Pain rating ~ time \* session \* (frequency change + frequency baseline) + (1 \| subject) |  |  |  |
| Conditional R<sup>2</sup> = .83 | Marginal R<sup>2</sup> = .07 |  |  |
| **Fixed effects and interactions** |  |  |  |
| Predictor | *F* (DF) | *η2* [95% CI] | *p* |
| Time | F (1, 17.9) = 21.54 | .11 [.05, 1] | <.001 |
| Session | F (1, 171.03) = 33.88 | .17 [.09, 1] | <.001 |
| Frequency change | F (1, 173.55) = .04 | <.01 [0, 1] | .85 |
| Frequency baseline | F (1, 188.68) = 3.28 | .02 [0, 1] | .07 |
| Time: session | F (1, 17.9) = 1.61 | <.01 [0, 1] | .21 |
| Time: frequency change | F (1, 171.34) = .11 | <.01 [0, 1] | .74 |
| Time: frequency baseline | F (1, 171.2) = .13 | <.01 [0, 1] | .72 |
| Session: frequency change | F (1, 172.25) = .2 | <.01 [0, 1] | .66 |
| Session: frequency baseline | F (1, 172.71) = .26 | <.01 [0, 1] | .61 |
| Time: session: frequency change | F (1, 171.37) = .63 | <.01 [0, 1] | .43 |
| Time: session: frequency baseline | F (1, 17.98) = .78 | <.01 [0, 1] | .38 |

For the α-band power of components identified through K-means clustering (Table 5) a significant effect of session was present (*F* (1, 147) = 1.16, *p* = .002) primarily driven by a significant pairwise comparison at 15 minutes post-HFS (*Std. effect* = .71 [.03, 1.39], *p*<.001), alongside an interaction with sex (*F* (1, 148) = 34.31, *p* < .001). This demonstrates that variations in α-band power across sessions were larger for female participants (*F* (1, 148) = 32, *p* < .001) when compared to male participants (*F* (1, 145) = 5.19, *p* = .02).

**Table 5.** K-means clustering α-band mixed-effects modelling. The linear mixed effects modelling results for k-means clustering α-band data are displayed for a) peak amplitude (power) b) PAF, and c-d) their relationships with MPS. = High-frequency stimulation. N = 27, (female = 19).

| **a) Model assessment: Power**<br>Amplitude ~ time \* session \* (age + sex) + (1 \| subject) |  |  |  |
|---|---|---|---|
| Conditional R<sup>2</sup> = .73 | Marginal R<sup>2</sup> = .11 |  |  |
| **Fixed effects and interactions** |  |  |  |
| Predictor | *F* (DF) | *η2* [95% CI] | *p* |
| Time | F (4, 141.12) = .52 | .01 [0, 1] | .72 |
| Session | F (1, 146.55) = 1.16 | .06 [.02, 1] | .002 |
| Age | F (1, 19.18) = .7 | .04 [0, 1] | .41 |
| Sex | F (1, 18.8) = .15 | <.01 [0, 1] | .7 |
| Time: session | F (4, 141.12) = .29 | <.01 [0, 1] | .88 |
| Time: age | F (4, 141.12) = .24 | <.01 [0, 1] | .92 |
| Time: sex | F (4, 141.12) = 1.41 | .04 [0, 1] | .23 |
| Session: age | F (1, 154.26) = .55 | <.01 [0, 1] | .46 |
| Session: sex | F (1, 148.19) = 34.31 | .19 [.1, 1] | <.001 |
| Time: session: age | F (4, 141.12) = .55 | .02 [0, 1] | .7 |
| Time: session: sex | F (4, 141.12) = .46 | .01 [0, 1] | .76 |
| **Between session comparisons: Session 2 – Session 1** |  |  |  |
| Minutes post-HFS | Estimate [95% CI] | Std. effect [95% CI] | *p* |
| -15 | .06 [-.28, .4] | .4 [-.28, 1.08] | .71 |
| 0 | .03 [-.2, .25] | .27 [-.41, .95] | .82 |
| 15 | .14 [.05, .24] | .71 [.03, 1.39] | <.001 |
| 30 | .1 [-.12, .31] | .54 [-.14, 1.22] | .37 |
| 45 | .13 [-.34, .6] | .68 [0, 1.36] | .57 |
| **Session 1 comparisons to baseline: Post-HFS – Baseline** |  |  |  |
| Minutes post-HFS | Estimate [95% CI] | Std. effect [95% CI] | *p* |
| 0 | .01 [-.3, .32] | .06 [-.62, .73] | 1 |
| 15 | -.12 [-.52, .28] | -.43 [-1.11, .25] | .83 |
| 30 | -.06 [-.37, .24] | -.24 [-.92, .43] | .92 |
| 45 | -.04 [-.51, .44] | -.15 [-.82, .53] | .99 |
| **Session 2 comparisons to baseline: Post-HFS – Baseline** |  |  |  |
| Minutes post-HFS | Estimate [95% CI] | Std. effect [95% CI] | *p* |
| 0 | -.03 [-.57, .52] | -.08 [-.74, .57] | 1 |
| 15 | -.04 [-.53, .46] | -.12 [-.78, .53] | .99 |
| 30 | -.03 [-.45, .39] | -.1 [-.76, .55] | .99 |
| 45 | .03 [-.5, .56] | .13 [-.53, .78] | 1 |
| **b) Model assessment: PAF**<br>Frequency ~ time \* session \* (age + sex) + (1 \| subject) |  |  |  |
| Conditional R<sup>2</sup> = .53 | Marginal R<sup>2</sup> = .18 |  |  |
| **Fixed effects and interactions** |  |  |  |
| Predictor | *F* (DF) | *η2* [95% CI] | *p* |
| Time | F (4, 141.11) = .23 | <.01 [0, 1] | .92 |
| Session | F (1, 152.36) = 4.76 | .03 [0, 1] | .03 |
| Age | F (1, 19.3) = 1.36 | .07 [0, 1] | .26 |
| Sex | F (1, 18.31) = 2.85 | .13 [0, 1] | .11 |
| Time: session | F (4, 141.11) = .33 | <.01 [0, 1] | .86 |
| Time: age | F (4, 141.11) = .21 | <.01 [0, 1] | .93 |
| Time: sex | F (4, 141.11) = 1.1 | .03 [0, 1] | .36 |
| Session: age | F (1, 159.97) = 6.17 | .04 [0, 1] | .01 |
| Session: sex | F (1, 154.51) = 5.92 | .04 [0, 1] | .02 |
| Time: session: age | F (4, 141.11) = 2.78 | .07 [0, 1] | .03 |
| Time: session: sex | F (4, 141.11) = .54 | .02 [0, 1] | .71 |
| **Between session comparisons: Session 2 – Session 1** |  |  |  |
| Minutes post-HFS | Estimate [95% CI] | Std. effect [95% CI] | *p* |
| -15 | -.46 [-1.49, .58] | -.52 [-1.2, .16] | .38 |
| 0 | -.4 [-.56, -.23] | -.47 [-1.14, .21] | <.001 |
| 15 | -.1 [-.98, .77] | -.07 [-.75, .6] | .82 |
| 30 | -.15 [-.74, .45] | -.15 [-.83, .52] | .62 |
| 45 | -.44 [-1.01, .12] | -.48 [-1.16, .2] | .12 |
| **Session 1 comparisons to baseline: Post-HFS – Baseline** |  |  |  |
| Minutes post-HFS | Estimate [95% CI] | Std. effect [95% CI] | *p* |
| 0 | .01 [-.4, .43] | .05 [-.62, .73] | 1 |
| 15 | -.26 [-.66, .15] | -.31 [-.98, .37] | .32 |
| 30 | -.09 [-.8, .61] | -.1 [-.78, .58] | .97 |
| 45 | -.09 [-.79, .61] | -.1 [-.78, .58] | .97 |
| **Session 2 comparisons to baseline: Post-HFS – Baseline** |  |  |  |
| Minutes post-HFS | Estimate [95% CI] | Std. effect [95% CI] | *P* |
| 0 | .08 [-1.26, 1.41] | .1 [-.55, .76] | 1 |
| 15 | .1 [-1.3, 1.5] | .14 [-.52, .79] | .99 |
| 30 | .22 [-1.07, 1.5] | .26 [-.39, .92] | .95 |
| 45 | -.08 [-1.37, 1.22] | -.07 [-.72, .59] | 1 |
| **c) Model assessment: Power influence on pain ratings**<br>Pain rating ~ time \* session \* (amplitude change + amplitude baseline) + (1 \| subject) |  |  |  |
| Conditional R<sup>2</sup> = .83 | Marginal R<sup>2</sup> = .07 |  |  |
| **Fixed effects and interactions** |  |  |  |
| Predictor | *F* (DF) | *η2* [95% CI] | *p* |
| Time | F (1, 167.62) = 3.85 | .16 [.08, 1] | <.001 |
| Session | F (1, 170.92) = 2.32 | .11 [.04, 1] | <.001 |
| Power change | F (1, 171.25) = .21 | <.01 [0, 1] | .65 |
| Power baseline | F (1, 185.69) = 8.82 | .05 [.01, 1] | <.001 |
| Time: session | F (1, 167.68) = .86 | <.01 [0, 1] | .35 |
| Time: power change | F (1, 168.14) = .46 | <.01 [0, 1] | .5 |
| Time: power baseline | F (1, 167.76) = .69 | <.01 [0, 1] | .41 |
| Session: power change | F (1, 169.87) = .9 | <.01 [0, 1] | .34 |
| Session: power baseline | F (1, 173.47) = 3.91 | .02 [0, 1] | .05 |
| Time: session: power change | F (1, 168.08) = .47 | <.01 [0, 1] | .49 |
| Time: session: power baseline | F (1, 167.68) = .47 | <.01 [0, 1] | .49 |
| **d) Model assessment: PAF influence on pain ratings**<br>Pain rating ~ time \* session \* (frequency change + frequency baseline) + (1 \| subject) |  |  |  |
| Conditional R<sup>2</sup> = .83 | Marginal R<sup>2</sup> = .1 |  |  |
| **Fixed effects and interactions** |  |  |  |
| Predictor | *F* (DF) | *η2* [95% CI] | *p* |
| Time | F (1, 167.87) = 34.65 | .17 [.09, 1] | <.001 |
| Session | F (1, 172.67) = 24 | .12 [.06, 1] | <.001 |
| Frequency change | F (1, 174.08) = .78 | <.01 [0, 1] | .38 |
| Frequency baseline | F (1, 186.57) = .08 | <.01 [0, 1] | .78 |
| Time: session | F (1, 167.89) = .8 | <.01 [0, 1] | .37 |
| Time: frequency change | F (1, 168.45) = .04 | <.01 [0, 1] | .85 |
| Time: frequency baseline | F (1, 168.11) = .01 | <.01 [0, 1] | .94 |
| Session: frequency change | F (1, 172.72) = .07 | <.01 [0, 1] | .79 |
| Session: frequency baseline | F (1, 172.7) = .36 | <.01 [0, 1] | .55 |
| Time: session: frequency change | F (1, 168.42) = .33 | <.01 [0, 1] | .56 |
| Time: session: frequency baseline | F (1, 167.99) = 2.35 | .01 [0, 1] | .13 |

For the PAF derived from K-means clustered components, a variable change between sessions was again evident (*F* (1, 152) = 4.76, *p* = .03) with a significant pairwise comparison immediately following HFS induction (*Std. effect* = -.47 [−1.14, .21], *p* < .001). Furthermore, this was mediated by both age (*F* (1, 160) = 6.17, *p* = .01) and sex (*F* (1, 155) = 5.92, *p* = .02), alongside a three-way interaction between session, time, and age (*F* (4, 141) = 2.78, *p* = .03). This demonstrates that PAF varied across sessions to a lesser degree for male participants (*F* (1, 151) = .09, *p* = .77) compared to female participants (*F* (1, 154) = 8, *p* = .005), and that for older participants (> 28 years) larger differences between sessions were evident at later time points.

When considering source localised sensorimotor α-band power (Table 6), no significant main effect was evident (*F* (1, 118.7) = .4, *p* = .53) despite large visual differences at the grand average level (Figure 1e). Instead, this was driven by large inter-individual differences based upon age (*F* (1, 116) = 4.16, *p* = .04) and sex (*F* (1, 119) = 6.05, *p* = .01). The largest variance between sessions occurred for female participants (*F* (1, 117) = 7.84, *p* = .006) compared to male participants (*F* (1, 120) = 1.06, *p* = .31), and older participants (> 44 years) compared to younger participants. In contrast, no significant main effects, interactions, or pairwise comparisons were evident for PAF derived from source localisation.

**Table 6.** Source localisation α-band mixed-effects modelling. The linear mixed effects modelling results for source localisation α-band data are displayed for a) peak amplitude (power) b) PAF, and c-d) their relationships with MPS. = High-frequency stimulation. N = 27, (female = 19).

| **a) Model assessment: Power**<br>Amplitude ~ time \* session \* (age + sex) + (1 \| subject) |  |  |  |
|---|---|---|---|
| Conditional R<sup>2</sup> = .74 | Marginal R<sup>2</sup> = .08 |  |  |
| **Fixed effects and interactions** |  |  |  |
| Predictor | *F* (DF) | *η2* [95% CI] | *p* |
| Time | F (4, 112.63) = .73 | .03 [0, 1] | .57 |
| Session | F (1, 118.7) = .4 | <.01 [0, 1] | .53 |
| Age | F (1, 16.57) = .58 | .03 [0, 1] | .46 |
| Sex | F (1, 16.56) = .32 | .02 [0, 1] | .58 |
| Time: session | F (4, 112.63) = .27 | <.01 [0, 1] | .9 |
| Time: age | F (4, 112.63) = .43 | .02 [0, 1] | .78 |
| Time: sex | F (4, 112.63) = .11 | <.01 [0, 1] | .98 |
| Session: age | F (1, 115.95) = 4.16 | .03 [0, 1] | .04 |
| Session: sex | F (1, 118.56) = 6.05 | .05 [.01, 1] | .01 |
| Time: session: age | F (4, 112.63) = .45 | .02 [0, 1] | .77 |
| Time: session: sex | F (4, 112.63) = .26 | <.01 [0, 1] | .9 |
| **Between session comparisons: Session 2 – Session 1** |  |  |  |
| Minutes post-HFS | Estimate [95% CI] | Std. effect [95% CI] | *p* |
| -15 | -.02 [-.67, .63] | -.1 [-.86, .65] | .96 |
| 0 | 0 [-.11, .11] | 0 [-.76, .76] | 1 |
| 15 | .05 [-.58, .67] | .31 [-.45, 1.07] | .88 |
| 30 | .04 [-.07, .16] | .2 [-.55, .96] | .47 |
| 45 | .06 [-.21, .33] | .34 [-.42, 1.1] | .68 |
| **Session 1 comparisons to baseline: Post-HFS – Baseline** |  |  |  |
| Minutes post-HFS | Estimate [95% CI] | Std. effect [95% CI] | *p* |
| 0 | .04 [-.79, .87] | .27 [-.48, 1.02] | 1 |
| 15 | 0 [-.78, .78] | 0 [-.75, .75] | 1 |
| 30 | .04 [-.8, .87] | .26 [-.49, 1.01] | 1 |
| 45 | .02 [-.73, .77] | .14 [-.61, .88] | 1 |
| **Session 2 comparisons to baseline: Post-HFS – Baseline** |  |  |  |
| Minutes post-HFS | Estimate [95% CI] | Std. effect [95% CI] | *p* |
| 0 | .06 [-.1, .21] | .38 [-.34, 1.09] | .71 |
| 15 | .06 [-.14, .26] | .41 [-.31, 1.12] | .8 |
| 30 | .1 [-.08, .27] | .57 [-.15, 1.29] | .42 |
| 45 | .1 [-.08, .27] | .58 [-.14, 1.3] | .42 |
| **b) Model assessment: PAF**<br>Frequency ~ time \* session \* (age + sex) + (1 \| subject) |  |  |  |
| Conditional R<sup>2</sup> = .67 | Marginal R<sup>2</sup> = .2 |  |  |
| **Fixed effects and interactions** |  |  |  |
| Predictor | *F* (DF) | *η2* [95% CI] | *p* |
| Time | F (4, 113.31) = .55 | .02 [0, 1] | .7 |
| Session | F (1, 122.24) = .57 | <.01 [0, 1] | .45 |
| Age | F (1, 17.15) = 2.01 | .1 [0, 1] | .17 |
| Sex | F (1, 17.16) = 3.81 | .18 [0, 1] | .07 |
| Time: session | F (4, 113.31) = .51 | .02 [0, 1] | .73 |
| Time: age | F (4, 113.31) = 1.45 | .05 [0, 1] | .22 |
| Time: sex | F (4, 113.31) = 1.21 | .04 [0, 1] | .31 |
| Session: age | F (1, 118.4) = .28 | <.01 [0, 1] | .6 |
| Session: sex | F (1, 122.06) = .1 | <.01 [0, 1] | .75 |
| Time: session: age | F (4, 113.31) = 2.43 | .08 [0, 1] | >.05 |
| Time: session: sex | F (4, 113.31) = .78 | .03 [0, 1] | .54 |
| **Between session comparisons: Session 2 – Session 1** |  |  |  |
| Minutes post-HFS | Estimate [95% CI] | Std. effect [95% CI] | *p* |
| -15 | .12 [-.34, .57] | .27 [-.49, 1.02] | .62 |
| 0 | -.06 [-.61, .48] | -.18 [-.93, .58] | .82 |
| 15 | .21 [-.25, .67] | .44 [-.32, 1.2] | .37 |
| 30 | -.1 [-.88, .68] | -.19 [-.95, .57] | .79 |
| 45 | .15 [-.68, .99] | .35 [-.4, 1.11] | .71 |
| **Session 1 comparisons to baseline: Post-HFS – Baseline** |  |  |  |
| Minutes post-HFS | Estimate [95% CI] | Std. effect [95% CI] | *p* |
| 0 | .04 [-.24, .32] | .13 [-.62, .88] | .97 |
| 15 | -.09 [-.4, .22] | -.17 [-.92, .58] | .84 |
| 30 | -.06 [-.34, .22] | -.13 [-.88, .62] | .93 |
| 45 | -.07 [-1.12, .99] | -.14 [-.89, .61] | 1 |
| **Session 2 comparisons to baseline: Post-HFS – Baseline** |  |  |  |
| Minutes post-HFS | Estimate [95% CI] | Std. effect [95% CI] | *P* |
| 0 | -.14 [-.79, .52] | -.31 [-1.03, .4] | .92 |
| 15 | 0 [-.64, .64] | 0 [-.71, .72] | 1 |
| 30 | -.27 [-1.12, .58] | -.58 [-1.3, .13] | .79 |
| 45 | -.03 [-.59, .53] | -.06 [-.77, .66] | 1 |
| **c) Model assessment: Power influence on pain ratings**<br>Pain rating ~ time \* session \* (amplitude change + amplitude baseline) + (1 \| subject) |  |  |  |
| Conditional R<sup>2</sup> = .86 | Marginal R<sup>2</sup> = .1 |  |  |
| **Fixed effects and interactions** |  |  |  |
| Predictor | *F* (DF) | *η2* [95% CI] | *p* |
| Time | F (1, 140.07) = 3.06 | .18 [.09, 1] | <.001 |
| Session | F (1, 144.43) = 15.89 | .1 [.04, 1] | <.001 |
| Power change | F (1, 147.3) = 2.03 | .01 [0, 1] | .16 |
| Power baseline | F (1, 157.09) = 2.18 | .01 [0, 1] | .14 |
| Time: session | F (1, 140.03) = 1.34 | <.01 [0, 1] | .25 |
| Time: power change | F (1, 140.05) = 1.7 | .01 [0, 1] | .19 |
| Time: power baseline | F (1, 139.9) = .43 | <.01 [0, 1] | .51 |
| Session: power change | F (1, 144.7) = .28 | <.01 [0, 1] | .59 |
| Session: power baseline | F (1, 151.21) = 6.97 | .04 [.01, 1] | .01 |
| Time: session: power change | F (1, 139.91) = .12 | <.01 [0, 1] | .73 |
| Time: session: power baseline | F (1, 139.81) = .53 | <.01 [0, 1] | .47 |
| **d) Model assessment: PAF influence on pain ratings**<br>Pain rating ~ time \* session \* (frequency change + frequency baseline) + (1 \| subject) |  |  |  |
| Conditional R<sup>2</sup> = .85 | Marginal R<sup>2</sup> = .07 |  |  |
| **Fixed effects and interactions** |  |  |  |
| Predictor | *F* (DF) | *η2* [95% CI] | *p* |
| Time | F (1, 139.86) = 29.6 | .17 [.09, 1] | <.001 |
| Session | F (1, 146.63) = 14.77 | .09 [.03, 1] | <.001 |
| Frequency change | F (1, 146.14) = .23 | <.01 [0, 1] | .63 |
| Frequency baseline | F (1, 157.72) = .7 | <.01 [0, 1] | .4 |
| Time: session | F (1, 139.86) = .32 | <.01 [0, 1] | .57 |
| Time: frequency change | F (1, 140.01) = 5.46 | .04 [0, 1] | .02 |
| Time: frequency baseline | F (1, 140.07) = .01 | <.01 [0, 1] | .94 |
| Session: frequency change | F (1, 141.57) = .02 | <.01 [0, 1] | .88 |
| Session: frequency baseline | F (1, 145.26) = 2.52 | .02 [0, 1] | .11 |
| Time: session: frequency change | F (1, 140.05) = 1.33 | <.01 [0, 1] | .25 |
| Time: session: frequency baseline | F (1, 139.94) = .1 | <.01 [0, 1] | .75 |
| **Session 1 relationships between frequency change and MPS:** |  |  |  |
| Minutes post-HFS | Trend [95% CI] | Std. effect [95% CI] | *p* |
| 0 | -.07 [-.14, .01] | -.29 [-.6, .03] | .08 |
| 15 | -.01 [-.07, .04] | -.06 [-.31, .18] | .62 |
| 30 | .04 [-.03, .11] | .16 [-.15, .48] | .31 |
| 45 | .09 [-.02, .2] | .39 [-.08, .85] | .1 |
| **Session 2 relationships between frequency change and MPS:** |  |  |  |
| Minutes post-HFS | Trend [95% CI] | Std. effect [95% CI] | *p* |
| 0 | -.03 [-.12, .07] | -.11 [-.52, .3] | .59 |
| 15 | -.01 [-.08, .06] | -.04 [-.33, .26] | .81 |
| 30 | .01 [-.05, .07] | .04 [-.22, .3] | .77 |
| 45 | .03 [-.05, .1] | .12 [-.22, .45] | .5 |

Together, these data suggest that component-level α-band features fluctuated widely between sessions in a manner that was driven by interindividual differences in age and sex. However, no significant changes over time were evident for either α-band power or PAF, suggesting that despite localising to sensorimotor activity, these methods failed to capture the HFS-induced changes evident at the sensor-level.

### Relationships between component-level resting-state EEG and MPS SHA

For each sensorimotor component identification method, mixed-effects modelling was used to investigate the relationship between α-band EEG features and MPS. For all analyses, significant main effects of time (*F* (1, 168) = 3.85, *p* < .001) and session (*F* (1, 171) = 2.32, *p* < .001) were apparent, evidencing the induction of SHA and differences in MPS between sessions as previously reported (Mugglestone, Ganis, et al., 2026). Therefore, for brevity, main effects for time and session are not reported separately for each case in the main text below.

For sensorimotor components identified through template matching, α-band power demonstrated a direct influence on MPS for both baseline amplitude (*F* (1, 183) = 6.55, *p* = .01) and change from baseline (*F* (1, 173) = 7.43, *p* = .01). This suggests that α-band power accurately reflected individual differences in MPS both at baseline and as MPS changed over time. Further investigation revealed that whilst the direction of relationship between MPS and α-band was consistent between sessions the magnitude of this effect was stronger during session 2 than session 1.

In contrast, template clustered sensorimotor component PAF demonstrated no influence on MPS for either baseline (*F* (1, 189) = 3.28, *p* = .07) nor change from baseline (*F* (1, 174) = .04, *p* = .85). Therefore, this methodology may be ineffective for identifying the PAF changes evident at the global sensor-level.

For k-means clustered components, whilst the influence of change from baseline power (*F* (1, 171) = .21, *p* = .65) was not significant, the effect of baseline power was (*F* (1, 186) = 8.82, *p* < .001), alongside an interaction between baseline power and changes in MPS across sessions (*F* (1, 173) = 3.91, *p* < .05). However, crucially, the interaction between baseline power and time was not significant (*F* (1, 168) = .69, *p* = .41). Therefore, whilst baseline sensorimotor α-band identified through k-means clustering may reflect inter-individual differences in pain ratings between sessions, this did not predict the development of SHA over time.

K-means clustered components also demonstrated no significant effects for baseline (*F* (1, 187) = .08, *p* = .78) or change from baseline (*F* (1, 174) = .78, *p* = .38) PAF on MPS.

For source localised sensorimotor components, there were no significant main effects of baseline (*F* (1, 157) = 2.18, *p* = .14) or change from baseline (*F* (1, 147) = 2.03, *p* = .16) power on MPS. However, a significant interaction between session and baseline power was evident (*F* (1, 151) = 6.97, *p* = .01), suggesting that interindividual differences in power influenced differences in MPS between sessions.

Alternatively, whilst no main effects were present for the influence of source localised PAF at baseline (*F* (1, 158) = .7, *p* = .4) or change from baseline (*F* (1, 146) < .01, *p* = .63), a significant interaction between change from baseline PAF and change in MPS over time was evident (*F* (1, 140) = 5.46, *p* = .02). This suggests that the predictive relationship between PAF change and MPS varied across time points. However, further investigation of estimated marginal trends revealed no significant individual relationship at any time point, with the significant interaction effect arising from a general shift in the direction of influence rather than a specifically important time point (see Table 6 for all estimates).

Together, these data suggest that sensorimotor component identification can capture distinctive elements of the global α-band signal. However, whilst the α-band power of components identified through template matching demonstrated significant influences on both inter- and intra-individual differences in MPS, given the lack of significant α-band power changes over time presented above, this measure is not in itself sensitive enough to derive changes to cortical activity related to SHA induction. K-means clustering also captured some of this relationship, likely driven by the overlapping components between the two methods, however, this was only sensitive to inter-individual changes in MPS between sessions. Furthermore, although components identified through source localisation also demonstrated a variable influence on α-band power and MPS, no significant relationship was evident at any time point. Additionally, no substantial SHA-related changes to PAF over time were evident for any of these component identification methods. Therefore, whilst changes to global PAF post-HFS tracked the development time course of SHA, the component identification methods failed to identify this.

## 5.2.5 Discussion

The present study investigated the temporal dynamics of HFS-induced changes to α-band power and PAF across repeated test sessions, alongside their inter- and intra-individual specificity for capturing variations in MPS throughout the SHA development time course. Global PAF demonstrated good test-retest reliability at baseline, and whist the strength of intra-individual changes from baseline varied across sessions they were found to reflect changes in MPS following HFS-conditioning. Furthermore, attempts to separate sensor-level data into local and global variance failed to identify a specific regional driver of the global post-HFS PAF slowing. This may suggest a widespread shift to thalamocortical communication rather than a specific change to somatosensory processing (Da Silva & Van Leeuwen, 1977; Da Silva et al., 1980; El Zghir et al., 2024; Halgren et al., 2019; Michalareas et al., 2016; Ploner et al., 2017).

In addition, methods of identifying and clustering sensorimotor components demonstrated low agreement and those identified through template matching did not consistently localise to pre- or post-central gyri. These component-level methods also failed to capture the global sensor-level PAF change, supporting sensitivity-related PAF slowing as a global phenomenon rather than a specific independent process. However, when assessed at residualised central electrodes and derived from sensorimotor components identified through template matching, a predictive relationship between α-band power and MPS was evident. Therefore, the present methods employed to derive sensorimotor-specific variance reveal that α-band power may be a useful marker of pain sensitivity depending upon how it is resolved from the global α-band signal.

The current study has demonstrated a significant relationship between changes in global PAF and MPS that was altered between sessions, indicating differences in the strength of relationship but not a systematic change in direction. This suggests that global PAF slowing was also sensitive to the previously reported (Mugglestone, Ganis, et al., 2026) alterations in the development time course of HFS-induced SHA, supporting slower PAF as a marker of increased pain sensitivity (Furman et al., 2018; Furman et al., 2020, 2021; Furman et al., 2019). However, the slowing of PAF post-HFS does not necessarily establish specificity to SHA and might instead reflect changes in fatigue or vigilance that also correspond to pain sensitivity, particularly as eyes-closed resting-state PAF is particularly susceptible to changes in alertness. Therefore, further replication is required, perhaps contrasting a time course of PAF change in the absence of HFS whilst retaining otherwise identical experimental conditions.

Despite these limitations, the present data align with previous experimental evidence that has demonstrated slowed PAF during ongoing tonic and capsaicin heat pain (Furman et al., 2021). The results of Furman et al. (2021) may also subtly link PAF to pain sensitivity given their use of capsaicin which can result in the concurrent development of SHA alongside the tonic sensation (Hughes et al., 2020; Magerl et al., 1998). However, the methods employed by Furman et al. (2021) were insufficient to separate whether changes to PAF were related to the intensity of ongoing tonic pain or arose from sensitivity changes as a result of the tonic stimulus, a discrepancy that is resolved by the present application of HFS, which induces SHA without an ongoing tonic pain sensation (Fawsitt-Jones et al., 2024; Klein et al., 2004; Klein et al., 2008; Lebrun, Lenoir, et al., 2025; Leone et al., 2025; Pfau et al., 2011; Quesada et al., 2021; van den Broeke et al., 2015).

Furthermore, the present data confirm that inter-individual differences in PAF prior to HFS-conditioning were not a trait predictor of HFS-induced SHA, supporting the conclusions of Lebrun, Ricci, et al. (2025) across repeated test sessions. However, this challenges the relationship between slower baseline PAF and the intensity of phasic and capsaicin heat pain as reported by Furman et al. (2020). This discrepancy may be due to the nociceptive stimuli applied, as pain resulting from ongoing tonic heat pain is distinct from acute mechanical pinpricks. Similarly, May et al. (2025) demonstrated that baseline PAF did not predict the perceived intensity of acute laser stimuli, validating this across a large cohort of healthy participants.

Whilst the slowing of PAF is evident across many long-term chronic conditions (Cavaleri et al., 2025; Cherin et al., 2026; Fauchon et al., 2022; Liberati, 2025; Mussigmann et al., 2022; Ploner et al., 2017), its mechanistic specificity has remained in question (Ahmad & Barkana, 2025; McLain et al., 2022; Mussigmann et al., 2022). Therefore, the present data provide clarification, demonstrating that PAF slowing can occur alongside the development of heightened pain sensitivity arising from spinally mediated plasticity, rather than functioning as a prior predictor of individual susceptibility to sensitivity development. Furthermore, as α-band oscillations represent postsynaptic communication within thalamocortical circuits (Da Silva & Van Leeuwen, 1977; Da Silva et al., 1980; El Zghir et al., 2024; Halgren et al., 2019; Michalareas et al., 2016; Ploner et al., 2017), alterations to these processes may be produced by the development of SHA.

Alternatively, whilst sensor-level PAF demonstrated moderate to good test-retest reliability at baseline across all electrode clusters, extremely poor reliability was evident when post-HFS change from baseline was assessed. Furthermore, despite significant variations between electrode groupings, no clear influence of PAF on MPS survived residualisation, nor were evident across component identification methods, supporting the hypothesis of Cherin et al. (2026) that the slowed PAF that is relevant to pain perception cannot be isolated to a single electrode cluster. Although Cherin et al. (2026) did not assess residualised data, they demonstrate that altered PAF relating to chronic pain is present across sensor-level groupings. Therefore, together with Cherin et al. (2026), the present data suggest that the identification of sensor-level PAF slowing at specific electrode groupings may risk producing artefactual regional effects that are essentially an expression of the global signal.

However, despite inconsistencies across sessions when assessed at the global sensor-level, residualised α-band power from the central electrode grouping was significantly predictive of differences in MPS between sessions, mirroring the reduced reported pain intensity at baseline previously reported (Mugglestone, Ganis, et al., 2026). Therefore, whilst slowed PAF may be a global phenomenon, inter-individual differences in α-band power that are relevant to pain perception may be specific to central electrodes. This regional specificity is further elucidated by the present component-level analyses which also demonstrated a significant relationship between α-band power and MPS at baseline for components identified through template matching. Which was further compounded by the continuing predictive relationship as MPS changed post-HFS. Therefore, sensorimotor α-band power may be sensitive to changes in acute pain perception. It is important to note that the authors are not implying that sensorimotor α-band power directly reflects pain perception, as the present data consider resting-state EEG, rather than the ERSP dynamics discussed in prior work (Mugglestone, Hughes., et al, 2026). Therefore, sensorimotor α-band power may capture something of the state of the nociceptive system prior to engagement, which was itself altered by HFS in a manner that corresponds to the degree and time course of SHA development.

Despite these complimentary analyses, this finding is surprising given largely inconsistent literature that has demonstrated variable relationships between α-band power and pain perception both across experimental conditions (Ahmad & Barkana, 2025; Bott et al., 2025; Ding et al., 2024; Furman et al., 2021; Lebrun, Ricci, et al., 2025), and within patient cohorts (Camargo et al., 2024; Fauchon et al., 2022; Mussigmann et al., 2022; Villafaina et al., 2019; Zebhauser et al., 2022), both in terms of direction (Ding et al., 2024; Furman et al., 2021; Villafaina et al., 2019) and presence (Camargo et al., 2024). This may be explained by differences in the methods used to resolve sensorimotor α-band power as no clear effect was evident when non-residualised sensor-level data were assessed, as is common in many studies. This suggests that even when assessed at specific electrode clusters, the global α-band signal may have overpowered regional-specific variance. Therefore, residualisation, component identification, or equivalent methods of isolating regional-specific variance may be critical for identifying relevant α-band power signals related to pain perception. Furthermore, the consideration of aperiodic features is crucial for accurately assessing α-band power (Donoghue, 2025; Donoghue et al., 2020), and is also often overlooked.

Additionally, the test-retest reliability of components identified through template matching was poor even at baseline and substantially reduced when change from baseline was assessed. Therefore, whilst template matching α-band power may presently demonstrate a feasible marker for indicating pain perception, this requires further replication and validation. Furthermore, agreement between component identification methods was low, which may evidence a distinction between components displaying a typical “sensorimotor” topographical projection (Cherin et al., 2026; Chowdhury et al., 2025) and those genuinely localising to sensorimotor gyri (Delorme & Makeig, 2004; Delorme et al., 2012; Desikan et al., 2006; Oostenveld et al., 2003; Oostenveld et al., 2011), although there was some overlap.

Finally, the present data demonstrate variability in α-band power and PAF across sensor-level electrode groupings attributable to inter-individual differences in age and sex. Greater variability across sessions was evident for older participants, which is consistent with previous research demonstrating that slower PAF and reduced α-band power is evident with age, likely contributing to greater measurement variability (Merkin et al., 2023). Furthermore, in all cases of significant difference between sexes, male participants demonstrated consistently slower PAF. However, when sex influenced the change in PAF between sessions or over time, no stable direction of influence was evident. This may be explained by differences between sexes in the underlying generators that contribute to the scalp PAF, as demonstrated through magnetoencephalography by (Fauchon et al., 2022). However, the present results do not offer a conclusive resolution.

In conclusion, these data have provided test-retest temporal dynamics of HFS-induced changes to resting-state EEG α-band power and PAF across repeated test sessions. Global PAF was reliable at baseline with a significant slowing post-HFS that differed in magnitude between sessions and was related to MPS. However, attempts to identify a specific regional driver of this global change failed, despite separating sensor-level data into local and global variance at multiple electrode groupings and the use of separate IC identification methods. Comparatively, residualised sensor-level analyses and ICs identified through template matching revealed a significant influence of central α-band power on MPS that may capture the state of the nociceptive system prior to engagement and retained predictive influence as SHA developed.

## Authorship Contributions

SM: Conceptualisation, funding acquisition, data collection, formal analysis, writing, reviewing and editing.

GG: Conceptualisation, funding acquisition, supervision, methodology, reviewing and editing.

SH: Conceptualisation, funding acquisition, supervision, methodology, reviewing and editing.

## Data availability statement

Data are available upon request.

## References

Ahmad, B., & Barkana, B. D. (2025). Pain and the Brain: A Systematic Review of Methods, EEG Biomarkers, Limitations, and Future Directions. Neurology International, 17(4), 46.

Association, W. M. (2013). World Medical Association Declaration of Helsinki: ethical principles for medical research involving human subjects. Jama, 310(20), 2191–2194.

Bannister, K., & Hughes, S. (2023). One size does not fit all: towards optimising the therapeutic potential of endogenous pain modulatory systems. Pain, 164(1), e5–e9.

Bates, D., Maechler, M., Bolker, B., Walker, S., Christensen, R. H. B., Singmann, H., Dai, B., Grothendieck, G., Green, P., & Bolker, M. B. (2015). Package ‘lme4’. convergence, 12(1), 2.

Biurrun Manresa, J., Kæseler Andersen, O., Mouraux, A., & van den Broeke, E. N. (2018). High frequency electrical stimulation induces a long-lasting enhancement of event-related potentials but does not change the perception elicited by intra-epidermal electrical stimuli delivered to the area of increased mechanical pinprick sensitivity. PLoS One, 13(9), e0203365.

Bland, J. M., & Altman, D. (1986). Statistical methods for assessing agreement between two methods of clinical measurement. The lancet, 327(8476), 307–310.

Bott, F. S., Zebhauser, P. T., Hohn, V. D., Turgut, Ö., May, E. S., Tiemann, L., Ávila, C. G., Heitmann, H., Nickel, M. M., & Day, M. A. (2025). Exploring electroencephalographic chronic pain biomarkers: a mega-analysis. EBioMedicine, 120.

Camargo, L., Pacheco-Barrios, K., Marques, L. M., Caumo, W., & Fregni, F. (2024). Adaptive and compensatory neural signatures in fibromyalgia: an analysis of resting-state and stimulus-evoked EEG oscillations. Biomedicines, 12(7), 1428.

Cavaleri, R., McLain, N. J., Heindel, M., Schrepf, A., Rodriguez, L. V., & Kutch, J. J. (2025). Peak alpha frequency is related to the degree of widespread pain, but not pain intensity or duration, among people with urologic chronic pelvic pain syndrome. Pain reports, 10(2), e1251.

Chang, C.-Y., Hsu, S.-H., Pion-Tonachini, L., & Jung, T.-P. (2018). Evaluation of artifact subspace reconstruction for automatic EEG artifact removal. 2018 40th annual international conference of the IEEE engineering in medicine and biology society (EMBC),

Cherin, J., McLain, N., Schrepf, A., Rodriguez, L. V., & Kutch, J. J. (2026). Association of widespread pain and peak alpha frequency cannot be isolated to a single network. Pain reports, 11(3), e1436.

Chhapola, V., Kanwal, S. K., & Brar, R. (2015). Reporting standards for Bland–Altman agreement analysis in laboratory research: a cross-sectional survey of current practice. Annals of Clinical Biochemistry, 52(3), 382–386.

Chowdhury, N. S., Bi, C., Furman, A. J., Chiang, A. K., Skippen, P., Si, E., Millard, S. K., Margerison, S. M., Spies, D., & Keaser, M. L. (2025). Predicting individual pain sensitivity using a novel cortical biomarker signature. JAMA neurology, 82(3), 237–246.

Da Silva, F. L., & Van Leeuwen, W. S. (1977). The cortical source of the alpha rhythm. Neuroscience letters, 6(2-3), 237–241.

Da Silva, F. L., Vos, J., Mooibroek, J., & Van Rotterdam, A. (1980). Relative contributions of intracortical and thalamo-cortical processes in the generation of alpha rhythms, revealed by partial coherence analysis. Electroencephalography and clinical neurophysiology, 50(5-6), 449–456.

Darrow, D., Herman, A., Hattab, T., König, S., Hermosillo, R., Moser, J., Koirala, S., Grimsrud, G., Krimmel, S., & Nahas, Z. (2025). An action networks model for pain reveals cortical neuromodulation targets.

Datta, D. (2017). blandr: a Bland-Altman Method Comparison package for R. Zenodo. doi, 10.

De Vet, H. C., Terwee, C. B., Knol, D. L., & Bouter, L. M. (2006). When to use agreement versus reliability measures. Journal of clinical epidemiology, 59(10), 1033–1039.

Delorme, A. (2023). EEG is better left alone. Scientific reports, 13(1), 2372.

Delorme, A., & Makeig, S. (2004). EEGLAB: an open source toolbox for analysis of single-trial EEG dynamics including independent component analysis. Journal of neuroscience methods, 134(1), 9–21.

Delorme, A., & Makeig, S. (2023). This is no “ICA bug”: response to the article,“ICA’s bug: how ghost ICs emerge from effective rank deficiency caused by EEG electrode interpolation and incorrect re-referencing”. Frontiers in Neuroimaging, 2, 1331404.

Delorme, A., Palmer, J., Onton, J., Oostenveld, R., & Makeig, S. (2012). Independent EEG sources are dipolar. PLoS One, 7(2), e30135.

Desikan, R. S., Ségonne, F., Fischl, B., Quinn, B. T., Dickerson, B. C., Blacker, D., Buckner, R. L., Dale, A. M., Maguire, R. P., & Hyman, B. T. (2006). An automated labeling system for subdividing the human cerebral cortex on MRI scans into gyral based regions of interest. Neuroimage, 31(3), 968–980.

Ding, K., Arginteanu, T., Anderson White, M., Lovell, L., Thakor, N. V., & Doshi, T. (2024). Electroencephalographic power ratio and peak frequency difference associate with central sensitization in chronic pain. Journal of neural engineering, 21(6), 066035.

Donoghue, T. (2025). A systematic review of aperiodic neural activity in clinical investigations. European Journal of Neuroscience, 62(7), e70255.

Donoghue, T., Haller, M., Peterson, E. J., Varma, P., Sebastian, P., Gao, R., Noto, T., Lara, A. H., Wallis, J. D., & Knight, R. T. (2020). Parameterizing neural power spectra into periodic and aperiodic components. Nature neuroscience, 23(12), 1655–1665.

Dosenbach, N. U., Raichle, M. E., & Gordon, E. M. (2025). The brain’s action-mode network. Nature Reviews Neuroscience, 26(3), 158–168.

El Zghir, R. K., Gabay, N. C., & Robinson, P. (2024). Unified theory of alpha, mu, and tau rhythms via eigenmodes of brain activity. Frontiers in Computational Neuroscience, 18, 1335130.

Esarey, J., & Sumner, J. L. (2018). Marginal effects in interaction models: Determining and controlling the false positive rate. Comparative Political Studies, 51(9), 1144–1176.

Fauchon, C., Kim, J. A., El-Sayed, R., Osborne, N. R., Rogachov, A., Cheng, J. C., Hemington, K. S., Bosma, R. L., Dunkley, B. T., & Oh, J. (2022). Exploring sex differences in alpha brain activity as a potential neuromarker associated with neuropathic pain. Pain, 163(7), 1291–1302.

Fawsitt-Jones, H., Vollert, J., O’Daly, O., Williams, S. C., McMahon, S. B., Howard, M. A., & Hughes, S. W. (2024). Reliability of quantitative sensory testing in the assessment of somatosensory function after high-frequency stimulation–induced sensitisation of central nociceptive pathways. Pain, 165(4), 941–950.

Fuchs, M., Kastner, J., Wagner, M., Hawes, S., & Ebersole, J. S. (2002). A standardized boundary element method volume conductor model. Clinical Neurophysiology, 113(5), 702–712.

Furman, A. J., Meeker, T. J., Rietschel, J. C., Yoo, S., Muthulingam, J., Prokhorenko, M., Keaser, M. L., Goodman, R. N., Mazaheri, A., & Seminowicz, D. A. (2018). Cerebral peak alpha frequency predicts individual differences in pain sensitivity. NeuroImage, 167, 203–210.

Furman, A. J., Prokhorenko, M., Keaser, M. L., Zhang, J., Chen, S., Mazaheri, A., & Seminowicz, D. A. (2020). Sensorimotor peak alpha frequency is a reliable biomarker of prolonged pain sensitivity. Cerebral cortex, 30(12), 6069–6082.

Furman, A. J., Prokhorenko, M., Keaser, M. L., Zhang, J., Chen, S., Mazaheri, A., & Seminowicz, D. A. (2021). Prolonged pain reliably slows peak alpha frequency by reducing fast alpha power. bioRxiv, 2021.2007. 2022.453260.

Furman, A. J., Thapa, T., Summers, S. J., Cavaleri, R., Fogarty, J. S., Steiner, G. Z., Schabrun, S. M., & Seminowicz, D. A. (2019). Cerebral peak alpha frequency reflects average pain severity in a human model of sustained, musculoskeletal pain. Journal of neurophysiology, 122(4), 1784–1793.

Gao, R., Peterson, E. J., & Voytek, B. (2017). Inferring synaptic excitation/inhibition balance from field potentials. NeuroImage, 158, 70–78.

González-Villar, A. J., Samartin-Veiga, N., Arias, M., & Carrillo-de-la-Peña, M. T. (2017). Increased neural noise and impaired brain synchronization in fibromyalgia patients during cognitive interference. Scientific reports, 7(1), 5841.

Gordon, E. M., Chauvin, R. J., Van, A. N., Rajesh, A., Nielsen, A., Newbold, D. J., Lynch, C. J., Seider, N. A., Krimmel, S. R., & Scheidter, K. M. (2023). A somato-cognitive action network alternates with effector regions in motor cortex. Nature, 617(7960), 351–359.

Gousset, S., Torta, D. M., Mouraux, A., Lambert, J., & van den Broeke, E. (2023). Pinprick-induced gamma-band oscillations are not a useful electrophysiological marker of pinprick hypersensitivity in humans. Clinical Neurophysiology, 153, 102–110.

Gyurkovics, M., Clements, G. M., Low, K. A., Fabiani, M., & Gratton, G. (2021). The impact of 1/f activity and baseline correction on the results and interpretation of time-frequency analyses of EEG/MEG data: A cautionary tale. NeuroImage, 237, 118192.

Haegens, S., Cousijn, H., Wallis, G., Harrison, P. J., & Nobre, A. C. (2014). Inter-and intra-individual variability in alpha peak frequency. NeuroImage, 92, 46–55.

Halgren, M., Ulbert, I., Bastuji, H., Fabó, D., Erőss, L., Rey, M., Devinsky, O., Doyle, W. K., Mak-McCully, R., & Halgren, E. (2019). The generation and propagation of the human alpha rhythm. Proceedings of the National Academy of Sciences, 116(47), 23772–23782.

Hamaker, E. L., & Muthén, B. (2020). The fixed versus random effects debate and how it relates to centering in multilevel modeling. Psychological methods, 25(3), 365.

Han, Q., Wang, H., Lu, X., Li, Y., Guo, Y., Zhao, X., Feng, Y., & Hu, L. (2025). Preoperative resting-state electrophysiological signals predict acute but not chronic postoperative pain. European Journal of Pain, 29(1), e4757.

Handiru, V. S., Suviseshamuthu, E. S., Yue, G., & Allexandre, D. (2024). Robust k-means-based Clustering of Independent Components Estimated from the EEG data. 2024 46th Annual International Conference of the IEEE Engineering in Medicine and Biology Society (EMBC),

Hsu, C. W., Schofield, T., & Ganis, G. (2024). Anterior N2 enhancement is not a general electrophysiological index of concealed information. Psychophysiology, 61(10), e14633.

Hsu, S.-H., Pion-Tonachini, L., Palmer, J., Miyakoshi, M., Makeig, S., & Jung, T.-P. (2018). Modeling brain dynamic state changes with adaptive mixture independent component analysis. NeuroImage, 183, 47–61.

Hughes, S. W., Basra, M., Chan, C., Parr, C., Wong, F., Gomes, S., & Strutton, P. H. (2020). Capsaicin-induced changes in electrical pain perception threshold can be used to assess the magnitude of secondary hyperalgesia in humans. Pain Medicine, 21(11), 2830–2838.

Kim, H., Luo, J., Chu, S., Cannard, C., Hoffmann, S., & Miyakoshi, M. (2023). ICA’s bug: How ghost ICs emerge from effective rank deficiency caused by EEG electrode interpolation and incorrect re-referencing. Frontiers in Signal Processing, 3, 1064138.

Kim, H., Miyakoshi, M., Kim, Y., Stapornchaisit, S., Yoshimura, N., & Koike, Y. (2022). Electroencephalography reflects user satisfaction in controlling robot hand through electromyographic signals. Sensors, 23(1), 277.

Klein, T., Magerl, W., Hopf, H.-C., Sandkühler, J., & Treede, R.-D. (2004). Perceptual correlates of nociceptive long-term potentiation and long-term depression in humans. Journal of Neuroscience, 24(4), 964–971.

Klein, T., Stahn, S., Magerl, W., & Treede, R.-D. (2008). The role of heterosynaptic facilitation in long-term potentiation (LTP) of human pain sensation. Pain, 139(3), 507–519.

Koo, T. K., & Li, M. Y. (2016). A guideline of selecting and reporting intraclass correlation coefficients for reliability research. Journal of chiropractic medicine, 15(2), 155–163.

Kuznetsova, A., Brockhoff, P. B., & Christensen, R. H. B. (2015). Package ‘lmertest’. R package version, 2(0), 734.

Lebrun, L., Lenoir, C., Leone, C., van den Broeke, E. N., Caspani, O., Schilder, A., Pelz, B., Truini, A., Treede, R.-D., & Mouraux, A. (2025). Strength, extent and duration of secondary hyperalgesia induced by high-frequency electrical stimulation of the foot compared to the volar forearm of healthy human volunteers. PLoS One, 20(2), e0318934.

Lebrun, L., Ricci, G., Courtin, A., van den Broeke, E. N., Lenoir, C., & Mouraux, A. (2025). Interindividual variations in peak alpha frequency do not predict the magnitude or extent of secondary hyperalgesia induced by high-frequency stimulation. Journal of neurophysiology, 134(6), 2025.2006. 2002.657417.

Legrain, V., & Mouraux, A. (2013). Activating selectively and reliably nociceptive afferentswith concentric electrode stimulation: yes we can! Provided that low stimulus intensities are used! Clinical Neurophysiology, 124(2), 424–424.

Lenth, R., Piaskowski, J., Banfai, B., Bolker, B., Buerkner, P., Giné-Vázquez, L., Hervé, M., Jung, M., Love, J., Miguez, F., Riebl, H., & Singmann, H. (2025). Package ‘emmeans’. R package version, 2(0), 107.

Leone, C., Di Lionardo, A., Di Pietro, G., Di Stefano, G., Falco, P., Blockeel, A., Caspani, O., Garcia-Larrea, L., Mouraux, A., & Phillips, K. (2021). How different experimental models of secondary hyperalgesia change the nociceptive flexion reflex. Clinical Neurophysiology, 132(12), 2989–2995.

Leone, C. M., Lenoir, C., & van den Broeke, E. N. (2025). Assessing signs of central sensitization: A critical review of physiological measures in experimentally induced secondary hyperalgesia. European Journal of Pain, 29(3), e4733.

Liberati, G. (2025). Neural biomarkers of pain: defining perspectives and limitations of peak alpha frequency. Pain.

Long, J. A., & Long, M. (2021). Package ‘interactions’. Comprehensive, User-Friendly Toolkit for Probing Interactions. Available online at: https://cran.r-project.org/web/packages/interactions/interactions.pdf (assessed July 29, 2021).

Lüdecke, D., Ben-Shachar, M. S., Patil, I., Waggoner, P., & Makowski, D. (2021). performance: An R package for assessment, comparison and testing of statistical models. Journal of open source software, 6(60).

Magerl, W., Wilk, S. H., & Treede, R.-D. (1998). Secondary hyperalgesia and perceptual wind-up following intradermal injection of capsaicin in humans. Pain, 74(2-3), 257–268.

May, E. S., Tiemann, L., Ávila, C. G., Bott, F. S., Hohn, V. D., Gross, J., & Ploner, M. (2025). Assessing the predictive value of peak alpha frequency for the sensitivity to pain. Pain, 10.1097.

McLain, N. J., Yani, M. S., & Kutch, J. J. (2022). Analytic consistency and neural correlates of peak alpha frequency in the study of pain. Journal of neuroscience methods, 368, 109460.

Medina, S., & Hughes, S. W. (2025). Immersion in nature through virtual reality attenuates the development and spread of mechanical secondary hyperalgesia: a role for insulo-thalamic effective connectivity. Pain, 166(9), 2181–2193.

Merkin, A., Sghirripa, S., Graetz, L., Smith, A. E., Hordacre, B., Harris, R., Pitcher, J., Semmler, J., Rogasch, N. C., & Goldsworthy, M. (2023). Do age-related differences in aperiodic neural activity explain differences in resting EEG alpha? Neurobiology of Aging, 121, 78–87.

Michalareas, G., Vezoli, J., Van Pelt, S., Schoffelen, J.-M., Kennedy, H., & Fries, P. (2016). Alpha-beta and gamma rhythms subserve feedback and feedforward influences among human visual cortical areas. Neuron, 89(2), 384–397.

Miyakoshi, M., Gehrke, L., Gramann, K., Makeig, S., & Iversen, J. (2021). The AudioMaze: An EEG and motion capture study of human spatial navigation in sparse augmented reality. European Journal of Neuroscience, 54(12), 8283–8307.

Mugglestone, S., Ganis, G., & Hughes, S. W. (2026). Temporal changes in mechanical pin prick sensitivity following high frequency induced sensitisation of central nociceptive pathways: a test re-test reliability study. The journal of pain, 106328.

Mugglestone, S., Hughes, S., & Ganis, G. (2026). Nociceptive-evoked electroencephalography can predict inter-individual differences in high frequency induced sensitisation of central nociceptive pathways: A test-retest reliability study. bioRxiv.

Mussigmann, T., Bardel, B., & Lefaucheur, J.-P. (2022). Resting-state electroencephalography (EEG) biomarkers of chronic neuropathic pain. A systematic review. NeuroImage, 258, 119351.

Nir, R.-R., Sinai, A., Raz, E., Sprecher, E., & Yarnitsky, D. (2010). Pain assessment by continuous EEG: association between subjective perception of tonic pain and peak frequency of alpha oscillations during stimulation and at rest. Brain research, 1344, 77–86.

Oostenveld, R., Delorme, A., & Makeig, S. (2003). DIPFIT: equivalent dipole source localization of independent components. In.

Oostenveld, R., Fries, P., Maris, E., & Schoffelen, J.-M. (2011). FieldTrip: open source software for advanced analysis of MEG, EEG, and invasive electrophysiological data. Computational intelligence and neuroscience, 2011(1), 156869.

Palmer, J. A., Kreutz-Delgado, K., & Makeig, S. (2012). AMICA: An adaptive mixture of independent component analyzers with shared components. Swartz Center for Computatonal Neursoscience, University of California San Diego, Tech. Rep, 35.

Pérez-Elvira, R., Cordeiro, D., Juan, M. A., & Sánchez Cabaco, A. (2026). An Exploratory Age-Adjusted Model of Individual Alpha Peak Frequency in Healthy Older Adults: A FOOOF-Based Spectral Parametrization Study. European Journal of Investigation in Health, Psychology and Education, 16(8), 110.

Perinelli, A., & Ricci, L. (2025). Stationarity assessment of resting state condition via permutation entropy on EEG recordings. Scientific reports, 15(1), 698.

Peterson, R. A., & Peterson, M. R. A. (2020). Package ‘bestNormalize’. Normalizing transformation functions. R package version, 1.

Pfau, D. B., Klein, T., Putzer, D., Pogatzki-Zahn, E. M., Treede, R.-D., & Magerl, W. (2011). Analysis of hyperalgesia time courses in humans after painful electrical high-frequency stimulation identifies a possible transition from early to late LTP-like pain plasticity. PAIN®, 152(7), 1532–1539.

Ploner, M., Sorg, C., & Gross, J. (2017). Brain rhythms of pain. Trends in cognitive sciences, 21(2), 100–110.

Poulsen, A. H., van den Berg, B., Arguissain, F., Tigerholm, J., Buitenweg, J. R., Andersen, O. K., & Mørch, C. D. (2022). Novel surface electrode design for preferential activation of cutaneous nociceptors. Journal of neural engineering, 19(1), 016010.

Quesada, C., Kostenko, A., Ho, I., Leone, C., Nochi, Z., Stouffs, A., Wittayer, M., Caspani, O., Brix Finnerup, N., & Mouraux, A. (2021). Human surrogate models of central sensitization: A critical review and practical guide. European Journal of Pain, 25(7), 1389–1428.

R Core Team. (2023). R: A Language and Environment for Statistical Computing. In R Foundation for Statistical Computing, Vienna, Austria.

Revelle, W. (2018). psych: Procedures for personality and psychological research. In: Northwestern University Evanston, IL.

Rousseeuw, P. J. (1987). Silhouettes: a graphical aid to the interpretation and validation of cluster analysis. Journal of computational and applied mathematics, 20, 53–65.

Seeley, W. W. (2019). The salience network: a neural system for perceiving and responding to homeostatic demands. Journal of Neuroscience, 39(50), 9878–9882.

Seeley, W. W., Menon, V., Schatzberg, A. F., Keller, J., Glover, G. H., Kenna, H., Reiss, A. L., & Greicius, M. D. (2007). Dissociable intrinsic connectivity networks for salience processing and executive control. The Journal of neuroscience, 27(9), 2349–2356.

Tadel, F., Baillet, S., Mosher, J. C., Pantazis, D., & Leahy, R. M. (2011). Brainstorm: A user-friendly application for MEG/EEG analysis. Computational intelligence and neuroscience, 2011(1), 879716.

Thammasan, N., & Miyakoshi, M. (2020). Cross-frequency power-power coupling analysis: A useful cross-frequency measure to classify ICA-decomposed EEG. Sensors, 20(24), 7040.

Umeda, T., Isa, T., & Nishimura, Y. (2019). The somatosensory cortex receives information about motor output. Science advances, 5(7), eaaw5388.

van den Broeke, E. N., De Hemptinne, P., Mercken, M., Torta, D., Lambert, J., & Mouraux, A. (2020). Central sensitization of nociceptive pathways demonstrated by robot-controlled pinprick-evoked brain potentials. Clinical Neurophysiology, 131(10), 2491–2498.

van den Broeke, E. N., De Vries, B., Lambert, J., Torta, D. M., & Mouraux, A. (2017). Phase-locked and non-phase-locked EEG responses to pinprick stimulation before and after experimentally-induced secondary hyperalgesia. Clinical Neurophysiology, 128(8), 1445–1456.

Van den Broeke, E. N., Lambert, J., Huang, G., & Mouraux, A. (2016). Central sensitization of mechanical nociceptive pathways is associated with a long-lasting increase of pinprick-evoked brain potentials. Frontiers in human neuroscience, 10, 531.

van den Broeke, E. N., Mouraux, A., Groneberg, A. H., Pfau, D. B., Treede, R.-D., & Klein, T. (2015). Characterizing pinprick-evoked brain potentials before and after experimentally induced secondary hyperalgesia. Journal of neurophysiology, 114(5), 2672–2681.

Villafaina, S., Collado-Mateo, D., Fuentes-Garcia, J. P., Cano-Plasencia, R., & Gusi, N. (2019). Impact of Fibromyalgia on Alpha-2 EEG Power Spectrum in the Resting Condition: A Descriptive Correlational Study. BioMed Research International, 2019(1), 7851047.

Viola, F. C., Thorne, J., Edmonds, B., Schneider, T., Eichele, T., & Debener, S. (2009). Semi-automatic identification of independent components representing EEG artifact. Clinical Neurophysiology, 120(5), 868–877.

Zebhauser, P. T., Hohn, V. D., & Ploner, M. (2022). Resting-state electroencephalography and magnetoencephalography as biomarkers of chronic pain: a systematic review. Pain, 164(6), 1200.

