## Supplementary Materials for "Intra-individual changes to peak alpha frequency map to pain perception following high frequency sensitisation of central nociceptive pathways: A test-retest reliability study"

Supplementary Table 1

**Sensor-level spectral parameterisation model fit assessments**

| Electrode grouping | $R^2$ (sd) | Mean error (sd) | Mean number of peaks (sd) |
|---|---|---|---|
| Session 1 | | | |
| Global | 0.95 (0.04) | 0.01 (0.01) | 2.36 (0.34) |
| Occipital | 0.94 (0.33) | 0.01 (0.01) | 2.28 (0.50) |
| Parietal | 0.98 (0.19) | 0.01 (0.01) | 2.36 (0.57) |
| Central | 0.98 (0.02) | 0.01 (0.01) | 2.5 (0.45) |
| Frontal | 0.97 (0.03) | 0.01 (0.01) | 2.47 (0.44) |
| Session 2 | | | |
| Global | 0.95 (0.04) | 0.01 (0.01) | 2.37 (0.32) |
| Occipital | 0.94 (0.06) | 0.01 (0.01) | 2.37 (0.53) |
| Parietal | 0.98 (0.02) | 0.01 (0.01) | 2.41 (0.42) |
| Central | 0.97 (0.03) | 0.01 (0.01) | 2.44 (0.43) |
| Frontal | 0.97 (0.02) | 0.01 (0.01) | 2.40 (0.5) |

*Supplementary Table 1 Sensor-level spectral parameterisation model fit assessments*

$R^2$, error, and number of peaks were identified separately for each spectral parameterisation model resulting in values for each participant, session, time point, and electrode. These were averaged across participants, time points, and electrodes as applicable for each session separately. sd = Standard deviation.

Supplementary Table 2

**Component-level spectral parameterisation model fit assessments**

| Clustering method | $R^2$ (sd) | Mean error (sd) | Mean number of peaks (sd) |
|---|---|---|---|
| Session 1 | | | |
| Template matching | 0.95 (0.13) | 0.01 (0.01) | 2.29 (0.63) |
| K-means clustering | 0.862 (0.24) | 0.01 (0.01) | 2.09 (0.77) |
| Source localisation | 0.9 (0.18) | 0.01 (0.01) | 2.18 (0.68) |
| Session 2 | | | |
| Template matching | 0.97 (0.05) | 0.01 (0.01) | 2.3 (0.7) |
| K-means clustering | 0.93 (0.16) | 0.01 (0.01) | 2.24 (0.71) |
| Source localisation | 0.92 (0.15) | 0.01 (0.01) | 2.12 (0.73) |

*Supplementary Table 2 Component-level spectral parameterisation model fit assessments*

$R^2$, error, and number of peaks were identified separately for each spectral parameterisation model resulting in values for each participant, session, time point, and electrode. These were averaged across participants, time points, and electrodes as applicable for each session separately. sd = Standard deviation.

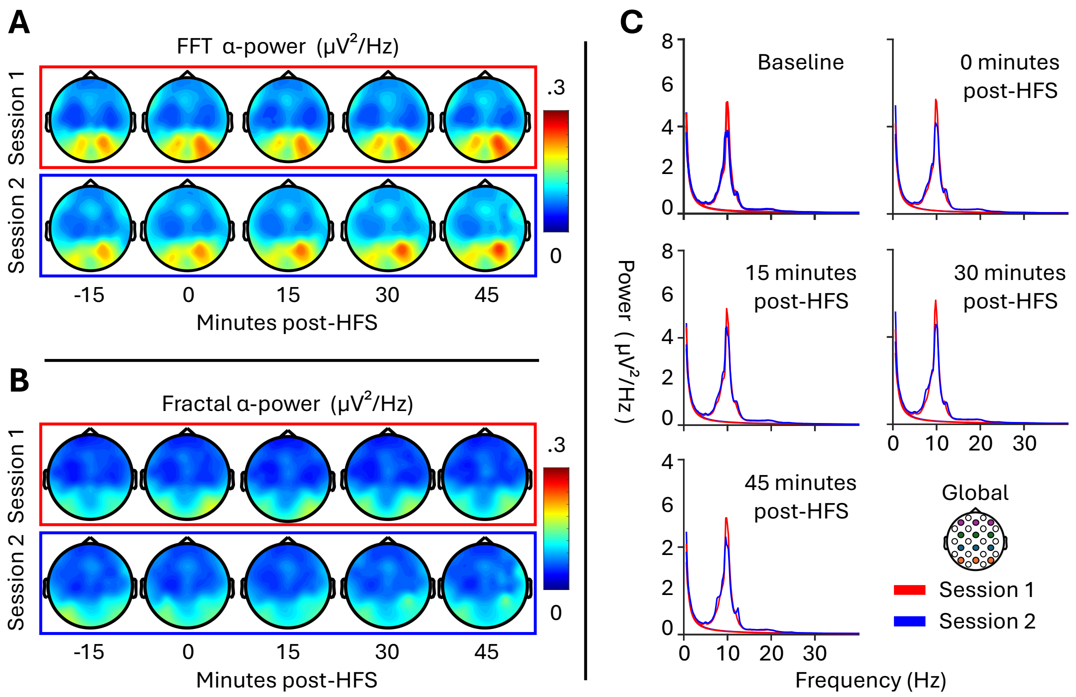


**Supplementary Figure 1 Resting-state EEG α-band data at the global average**

Global α-band topographies display a) FFT and b) Fractal power grand-averaged across all participants. The c) equivalent spectral frequency data represent the global average across all electrodes. Topographical plots were derived from a slightly wider 7 Hz to 13 Hz α-band to preserve data context across participants. FFT refers to the averaged spectral data following Fast-Fourier Transform. Fractal refers to the averaged estimated aperiodic component using Spectral Parametisation. HFS = High-frequency stimulation.

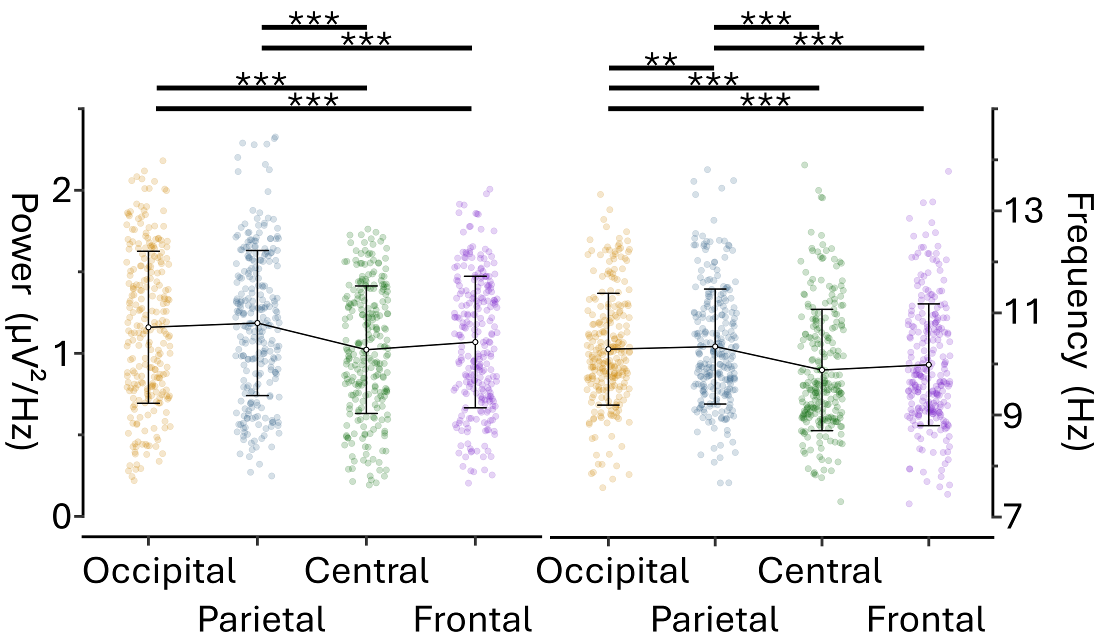


**Supplementary Figure 2 Differences in resting-state EEG α-band data between electrode groupings**

α-band peak power (left) and frequency (right) are displayed as individual datapoints across all time points and sessions for each individual. Black bars indicate group means and 1 standard deviation. Statistics were derived from linear mixed-effects modelling. ** indicates significance at alpha < .01. *** indicates significance at alpha < .001.

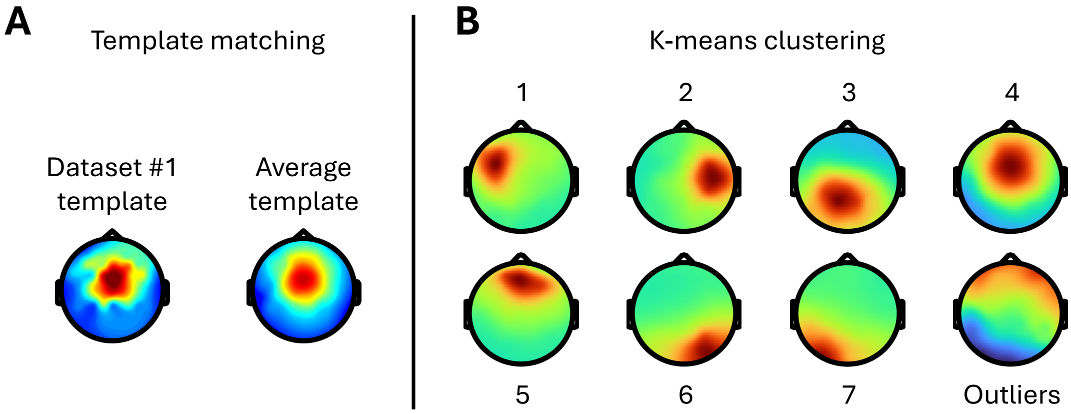


**Supplementary Figure 3 Resting-state EEG component selection templates and clusters**

Component topographies display a) the templates used for template matching sensorimotor component identification and b) the average of all components within each cluster following K-means clustering. Whilst there was a 93% carryover from the single-subject template to the group-average template, correlation to the average improved component selection with a mean correlation of .85. Of the 7 component clusters identified through K-means clustering, cluster 4 was selected as the sensorimotor cluster.

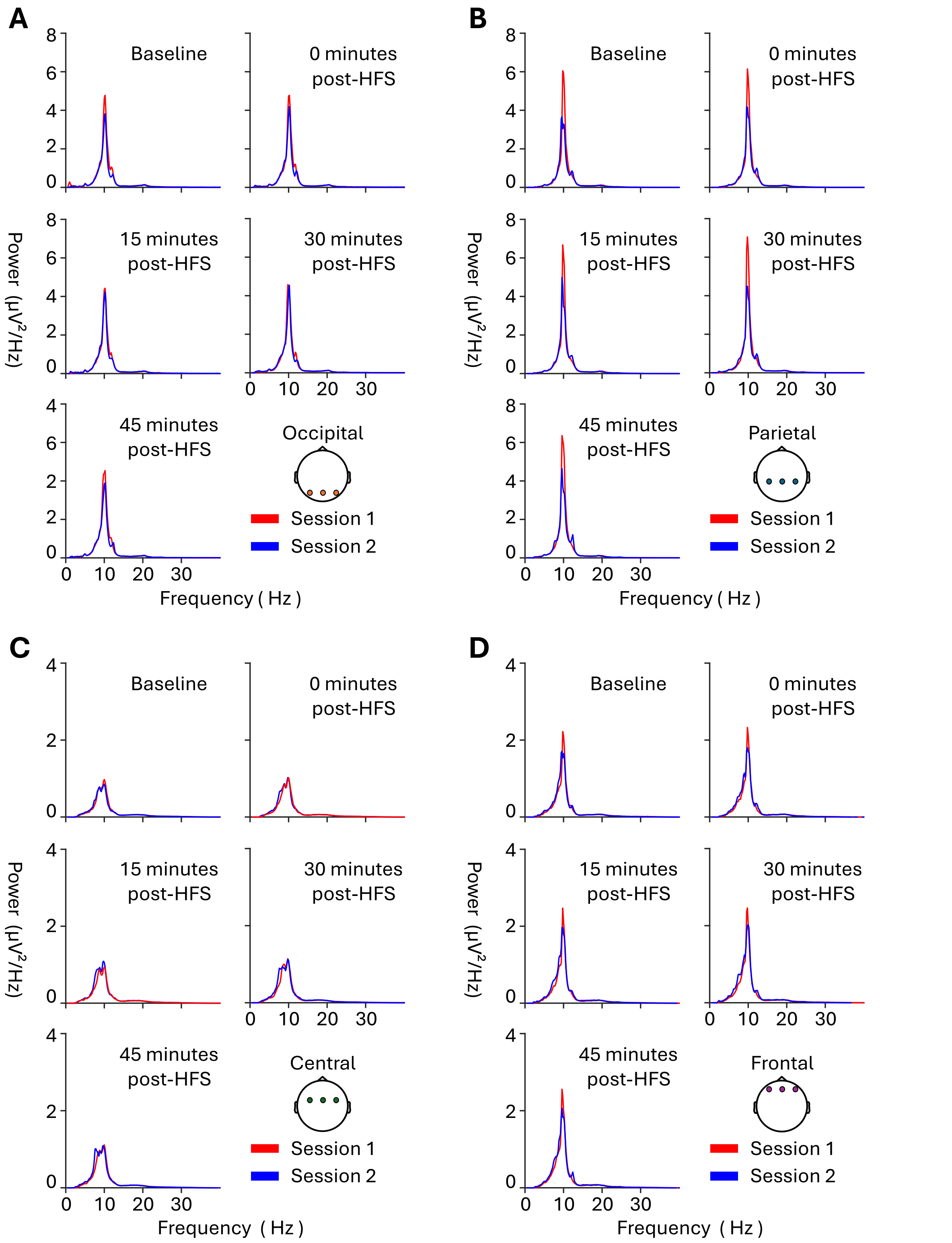


**Supplementary Figure 4 Resting-state EEG α-band data across electrode groupings**

Power spectrum plots display resting-state EEG α-band data grand-averaged across all participants for each time point and session at a) occipital electrodes, b) parietal electrodes, d) central electrodes, and e) frontal electrodes. PAF = Peak alpha frequency; HFS = High-frequency stimulation.

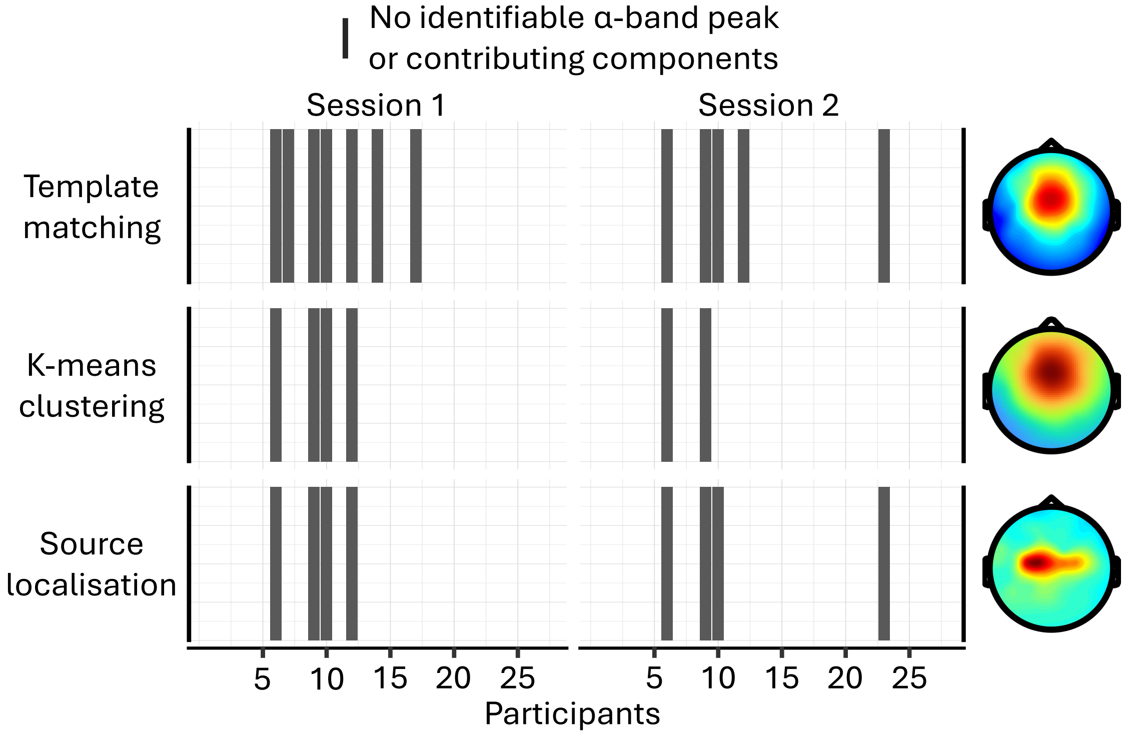


**Supplementary Figure 5 Proportion of datasets failing to contribute to component identification methods**

Each bar represents a dataset that either had no identifiable α-band peak (template matching and source localisation) or failed to contribute to the sensorimotor cluster (K-means clustering).
